# Behavioral-transcriptomic landscape of engineered T cells targeting human cancer organoids

**DOI:** 10.1101/2021.05.05.442764

**Authors:** Johanna F. Dekkers, Maria Alieva, Astrid Cleven, Farid Keramati, Peter Brazda, Heggert G. Rebel, Amber K.L. Wezenaar, Jens Puschhof, Maj-Britt Buchholz, Mario Barrera Román, Inez Johanna, Angelo D. Meringa, Domenico Fasci, Maarten H. Geurts, Hendrikus C.R. Ariese, Esmée J. van Vliet, Ravian L. van Ineveld, Effrosyni Karaiskaki, Oded Kopper, Yotam E. Bar-Ephraim, Kai Kretzschmar, Alexander M.M. Eggermont, Ellen J. Wehrens, Henk G. Stunnenberg, Hans Clevers, Jürgen Kuball, Zsolt Sebestyen, Anne C. Rios

## Abstract

Cellular immunotherapies are rapidly gaining clinical importance, yet predictive platforms for modeling their mode of action are lacking. Here, we developed a dynamic immuno-organoid 3D imaging-transcriptomics platform; BEHAV3D, to unravel the behavioral and underlying molecular mechanisms of solid tumor targeting. Applied to an emerging cancer metabolome-sensing immunotherapy: TEGs, we first demonstrate targeting of multiple breast cancer subtypes. Live-tracking of over 120,000 TEGs revealed a diverse behavioral landscape and identified a ‘super engager’ cluster with serial killing capability. Inference of single-cell behavior with transcriptomics identified the gene signature of ‘super engager’ killer TEGs, which contained 27 genes with no previously described T cell function. Furthermore, guided by a dynamic type 1 interferon (IFN-I) signaling module induced by high TEG-sensitive organoids, we show that IFN-I can prime resistant organoids for TEG-mediated killing. Thus, BEHAV3D characterizes behavioral-phenotypic heterogeneity of cellular immunotherapies and holds promise for improving solid tumor-targeting in a patient-specific manner.

## Introduction

Single-cell analyses are providing unprecedented opportunities to analyze the complexity of biological systems (van der Leun et al., 2020). However, they are restricted to providing a snapshot of cellular processes at a given timepoint. Yet living cells are highly dynamic, and their dynamic behavior shapes their function. Therefore, the development of technologies that address individual cell dynamics within a population is essential for understanding cellular behaviors and how these behaviors relate to function. Immune cells engineered to locate and kill tumor cells represent such dynamic cell populations with an increasing clinical importance (June and Sadelain, 2018). Successes of T cell therapies for hematological malignancies have sparked efforts to translate such approaches to solid tumors, including breast cancer (BC), but efficacy has so far been limited (Chen and Mellman, 2017). This poses a clear need for better understanding the mechanism of action of cellular therapies in order to optimize treatment design.

Because of challenges in identifying tumor-specific antigens for solid cancers (Schumacher et al., 2019), pan-tumor therapies that recognize metabolic alterations in cancer cells are being explored (Crowther et al., 2020). This includes an emerging therapy called TEGs, which are peripheral blood αβ T cells engineered to express a Vγ9/Vδ2 T cell receptor (TCR), comprising both CD4^+^ and CD8^+^ subsets (Gründer et al., 2012; Johanna et al., 2019; Marcu-Malina et al., 2011; Sebestyen et al., 2019; Vyborova et al., 2020). These hybrid cells have the ability to recognize cancer cells via the Vγ9/Vδ2 TCR that senses metabolic changes through the recently identified ligand butyrophilin 2A1 (BTN2A1) bound to BTN3A1 (Rigau et al., 2020; Vyborova et al., 2020). Yet, they maintain the high proliferation and memory capacity of conventional αβT cells (Marcu-Malina et al., 2011). TEGs are currently in clinical trials for various leukemia (Sebestyen et al., 2019), but their potential to target solid tumors remains unknown and should be adressed in adequate preclinical models.

There is a growing interest to use organoid technology to model immunotherapy function (Bar-Ephraim et al., 2019; Cattaneo et al., 2020; Neal et al., 2018; Schnalzger et al., 2019; Dijkstra et al., 2018). Patient-derived organoids (PDOs) provide reliable *in vitro* human cancer models that recapitulate important characteristics of the original tumor specimen (Tuveson et al.), allowing for the study of patient-specifc therapy responses (Ganesh et al., 2019; Ooft et al., 2019; Tiriac et al., 2018; Vlachogiannis et al., 2018; Yao et al., 2020). In addition, imaging has proven to be a powerful approach to characterize the spatial cellular organization and tissue dynamics in these 3D structures (Dekkers et al., 2019; van Ineveld et al., *in press*; Lukonin et al., 2020; 2019; Serra et al., 2019). Here, we aim to combine organoid and 3D imaging technology for the analysis of functional single cell behavior integrated with transcriptomic profiling to decipher and manipulate the solid tumor-targeting strategy of engineered immune cells (**Video S1**).

## Results

### 3D live-tracked TEG targeting efficacy

We devised a multispectral 3D image-based platform; BEHAV3D, to live-track efficacy and mode-of-action of cellular immunotherapy for ∼60 human cancer organoid cultures simultaneously (**Figures 1A-1C; Video S1**). Multiple real-time fluorescent dyes and nanobody technology were implemented for single acquisition 4-fluorophore spectral 3D imaging of T cell populations, organoids, and dead cells, allowing us to track single TEGs, individual organoids and their death-dynamics over 24 h (**Figures 1B, 1C, and S1A-S1C**). We detected a high variation of TEG-mediated killing efficacy in cultures derived from 14 BC patients (**Figure 1D; Table S1**), and different targeting kinetics over time (**Figures 1E, 1F, and S1D-S1F**), with percentages of dying PDOs ranging from near 0% (e.g. 34T) to 100% (e.g. 13T) (**Figure 1F**). Pearson correlation analysis between imaging data and a commonly used cell viability assay **(Figures S1G and S1H)**, or interferon gamma (IFN-γ) secretion measured by ELISA **(Figures S1I and S1J)**, confirmed robustness of our imaging quantification method. Among the 6 highest TEG-sensitive PDO cultures (above 50% dying organoids; **Figure 1F**), we noted cultures derived from primary BC of distinct subtypes (triple negative breast cancer (TNBC), human epidermal growth factor receptor 2 (HER2)^+^, estrogen receptor (ER)^+^, and ER^+^ progesterone receptor (PR)^+^HER2^+^) and from a metastasis derived from a HER2^+^ primary tumor (**Figures 1D and 1F**), supporting the potential of TEGs in targeting a broad spectrum of BCs. Importantly, TEGs control the growth of PDO-derived breast tumors (TNBC primary tumor and HER2^+^ metastasis) in mouse xenograft models (**Figure 1G**), showing efficacy of TEG for BC *in vivo*.

**Figure 1.**
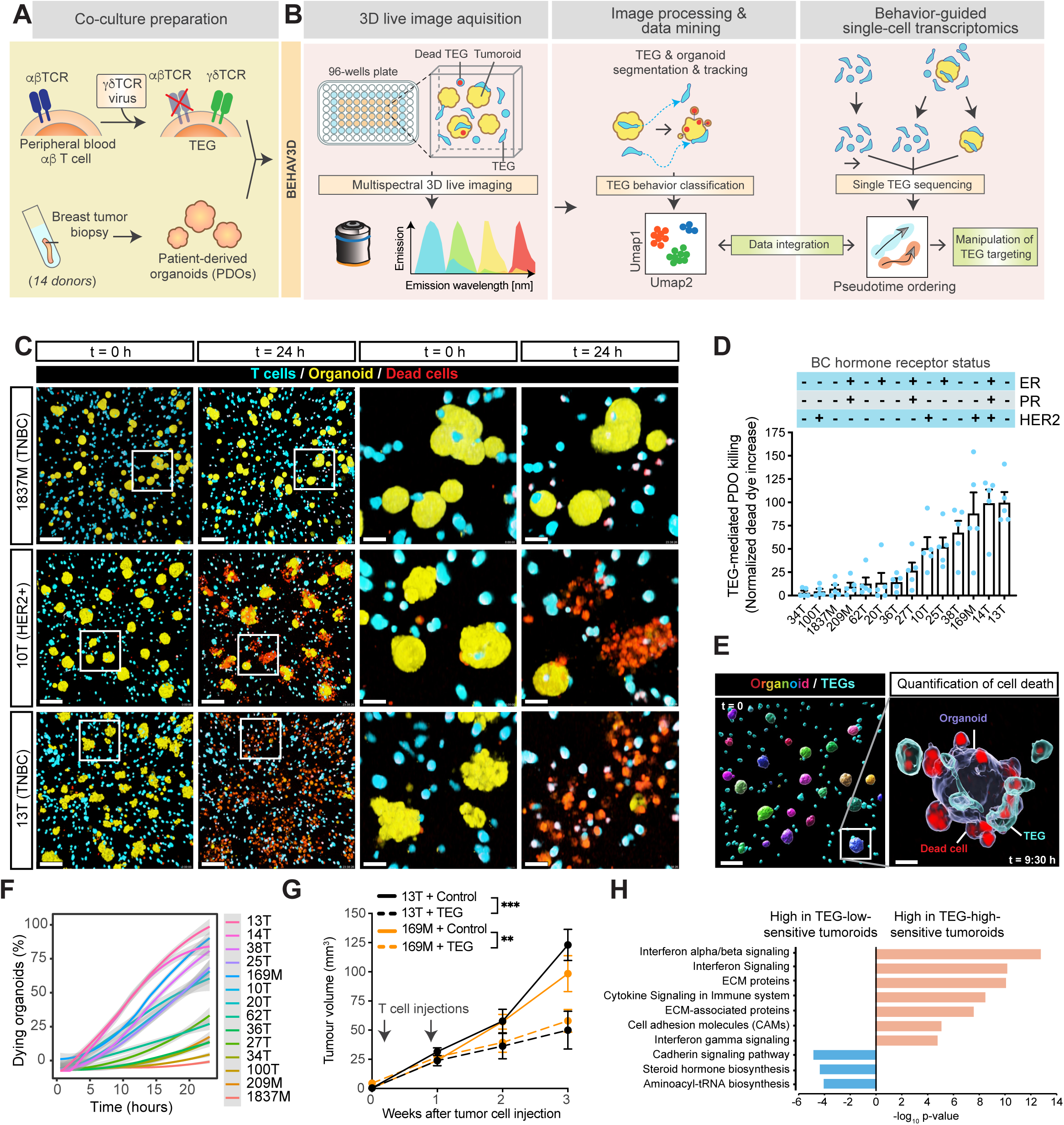
TEG efficacy across organoids of multiple breast cancer subtypes detected by multispectral 3D live imaging and *in vivo* TEG targeting. (**A**) Schematic representation of co-culture preparation. TEGs were generated by engineering peripheral blood αβ T cells to express a defined Vγ9/Vδ2 TCR via retroviral transduction. TEGs were then co-cultured with patient-derived breast organoids (PDOs). (**B**) Schematic representation of the BEHAV3D platform. Fluorescent dyes were combined to specifically label organoids (yellow), TEGs (blue) and dead cells (red). Co-cultures of organoids and TEGs were imaged in 96-well plates using spectral confocal microscopy in 3D, followed by segmentation and tracking of organoids and T cells, and subsequent behavior classification. TEGs of experimental conditions as indicated were sequenced and pseudotime ordering was used to integrate behavioral data. Identified targets were used to manipulate TEG targeting. (**C**) Representative 3D multispectral images of breast PDO cultures (yellow) that show low (1837M), intermediate (10T) and high (13T) killing by TEGs (blue) at the indicated time points of imaging. Dead cells depicted in red. Scale bars, 100 µm (left two columns) and 30 µm (right two columns). (**D**) Quantification of killing of organoids derived from 14 different BC patients upon 24 hr co-culture with TEGs by 3D live cell imaging. All data were corrected for control LM1 T cell responses. (n = 4 independent experiments; mean ± s.e.m.; TNBC = triple negative breast cancer; ER = estrogen receptor; PR = progesterone receptor). (**E**) Representative 3D multispectral images showing automated rendering of single organoids (confetti colors) and T cells (blue) (left image), and an enlarged section showing presence of dead cell dye (red) in a single organoid (transparent purple rendering) and TEGs (transparent blue rendering) at the indicated time of co-culture. Scale bars, 100 µm (left image) and 30 µm (right image). (**F**) Quantification of the percentage of dying single organoids (% of total) over time for each PDO co-cultured with TEGs (n = 4 independent experiments; mean ± s.e.m.). (**G**) Quantification of the volume of tumors overtime generated by subcutaneous transplantation of 13T (black lines) or 169M organoids (orange lines). Animals received 2 injections of either TEGs (dashed line) or control TEG011 cells (Control; solid line) at the indicated timepoints. (n ≥5 per condition; mean ± s.e.m.). Statistical analysis was performed by Two-Way ANOVA with repeated measures: 13T-TEG vs 13T-control p < 0.0001; 169M-TEG vs 169M-control p=0.0016. (**H**) Gene ontology enrichment analysis of differentially expressed genes between the six highest versus six lowest TEG-sensitive organoid cultures from d.

### PDO inflammatory features associate with TEG sensitivity

Bulk RNA sequencing of PDOs revealed differentially expressed genes (DEGs) between the 6 lowest versus the 6 highest TEG-sensitive PDO cultures (**Table S2**), related to upregulated cadherin signaling and steroid biosynthesis pathways in TEG-insensitive cultures, whereas cytokine signaling, as well as extracellular matrix (ECM) organization, correlated with high sensitivity to TEG therapy (**Figures 1H, and S1K-S1M**). The highest association was found between TEG killing and type 1 interferon (IFN-I) signaling genes, including MX1, IFIT1, OASL, and XAF1, which were highly expressed especially in the 2 highest TEG-sensitive PDO cultures; 14T and 13T (**Figures 1H, and S1M)**. Thus, PDOs maintain tumor-specific inflammatory features in culture, highlighting their utility for modeling cellular immunotherapy responses in a patient-specific manner.

### TEGs display a high diversity in behavior and killing potential

BEHAV3D implements single immune cell tracking in a 3D space over time and behavioral classification (**Figures 1B, and 2A; Video S1**), revealing that -when exposed to PDOs-TEGs could be separated into nine subpopulations with unique behavioral patterns (**Figures 2B-2D, and S2A, S2B**). Patterns varied from inactive behaviors (*dying*, *static* and *lazy*) to active motility (*slow scanner*, *medium scanner* and *super scanner*) and organoid engagement (*tickler*, *engager* and *super engager*), thus demonstrating a high level of behavioral heterogeneity. Having captured their behavioral single-cell landscape in this classifier (**Figures S3C-S3E)**, we next predicted TEG behavior when co-cultured with PDOs that showed varying TEG sensitivity (34T, 100T, 27T, 10T or 13T; **Figure 1F**), as well as an organoid culture derived from normal breast tissue, which only showed minimal death when cultured with TEGs (**Figure 2E**). A total of 123,296 TEGs were live-tracked to investigate how the organoid (inflammatory) profile shapes T cell behavior. For each PDO culture, TEGs displayed unique distributions of behavioral signatures (**Figure 2E**) and higher organoid killing associated with an increase in tumor engagement (*tickler*, *engager* and *super engager*), while *static, lazy* and *medium scanner* behavior decreased (**Figure 2F**). Correlation between single organoid dying dynamics and TEG engagement over time revealed that organoids contacted by *super engagers*, as compared to other organoid-engaging clusters, had the highest chance of being killed (**Figures 2G, and S2F).** This indicates that effective killing by TEGs relies on prolonged organoid contact, a main feature of *super engagers* (48 ± 8 min / hr; mean ± s.d.).

**Figure 2.**
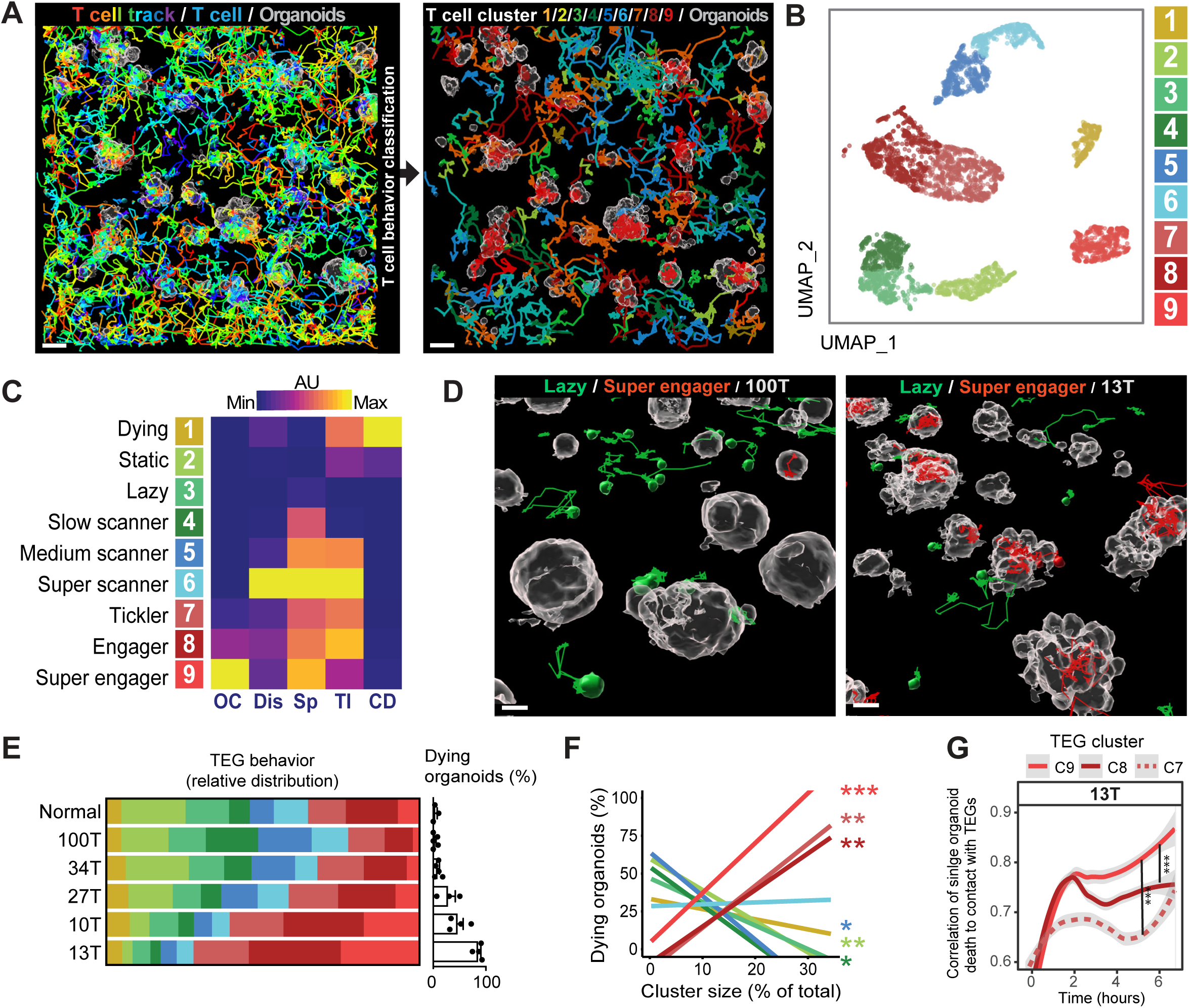
TEGs exposed to PDOs display high diversities in their behavior with distinct killing potential. (**A**) Representative image of automated tracking of each TEG (left image; 10 hrs tracks are rainbow-colored for time). Tracks were classified according to TEG behavior and back-projected in the image (right image; color-coded by cluster). Scale bars, 50 µm. (**B**) Umap plot showing nine color-coded clusters identified by unbiased multivariate timeseries dynamic time warping analysis. Each data point represents a T cell track of 3.3 hrs. Numbers refer to cluster names presented in (C). (**C**) Heatmap depicting relative values of T cell features indicated for each cluster, named according to their most distinct characteristics. AU: arbitrary units in respect to maximal and minimal values for each feature. (OC, organoid contact; Dis, square displacement; Sp, speed; TI, T cell interactions; CD, cell death) (**D**) 3D-rendered images of 100T (low-targeting; left image) and 13T (high-targeting; right image) organoids (grey) and TEGs with 3.3 hr tracks belonging to *lazy* (green) and *super engager* (red) clusters. Scale bars, 20 µm. (**E**) Behavioral cluster distribution of TEGs co-cultured with the indicated PDOs and a normal organoid culture (left plot), in relation to their killing capacity (right bar graph) represented as the percentage of dying organoids (% of total) (n ≥ 3 independent experiments; mean ± s.e.m.). X^2^ test; p = 1.132e-08. (**F**) Pearson correlation between behavior cluster size and the percentage of dying organoids represented in d. CL9 p=0.00006; CL8 p= 0.009; CL7 p=0.006; CL5 p=0.014; CL4 p=0.022; CL2 p=0.0019. (n ≥ 3 independent experiments; mean). (**G**) Change in correlation between 13T organoid death dynamics (measured as increase in dead cell dye) and cumulative contact with TEGs (from cluster (C)7-9). Data is represented as mean correlation per timepoint of all single organoids (n = 4 independent experiments). Linear mixed model fitting with each experimental replicate as a random effect: C9 vs C8 p= 5.19e-06; C9 vs C7 p < 2e-16.

### Serial killing capability of *super engager* CD8^+^ TEGs

We next linked behaviors to population phenotypes by first differentially labeling CD4^+^ and and CD8^+^ T cells **(Figures 3A, and S3A**). This revealed that prolonged organoid contact and *super engager* behavior was a preferred feature of CD8^+^ TEGs, whereas CD4^+^ TEGs showed a higher proportion of *lazy* cells, *slow scanners*, *medium scanners*, *super scanners*, and *ticklers* (**Figures 3A-3C)** characteristic of high movement and short organoid contact (**Figure 2C)**. Furthermore, long-term behavior classification and back-projection of cells classified in the live-tracked imaging dataset (**Figures S3B and S3C**), showed that single CD8^+^ TEGs, once engaged with an organoid, most often killed multiple cells consecutively (serial killing) (**Figures 3D-3G**), a preferred feature of engineered T cells(Cazaux et al., 2019; Halle et al., 2016; Weigelin et al.). In contrast, CD4^+^ TEGs often moved away after organoid engagement without killing, but occasionally targeted individual cells in different organoids (**Figures 3D-3F, and S3D**) thereby displaying slower killing rates (**Figure 3H**). Serial killing by *super engager* CD8^+^ TEGs was characterized by attachment to PDOs using a defined anchor point from where surrounding cells were killed via long protrusions, intercalating between epithelial cells and extending their initial size up to 5 times (**Figures 3E, and S3E and S3F**). Remarkably, single CD8^+^ TEGs were able to kill entire organoids (up to 18 cells in 11 hrs; **Figures 3E and 3G; Video S1**). This extent of serial killing and morphological plasticity of *super engager* CD8^+^ TEGs was uniquely revealed by the high spatiotemporal resolution character of BEHAV3D.

**Figure 3.**
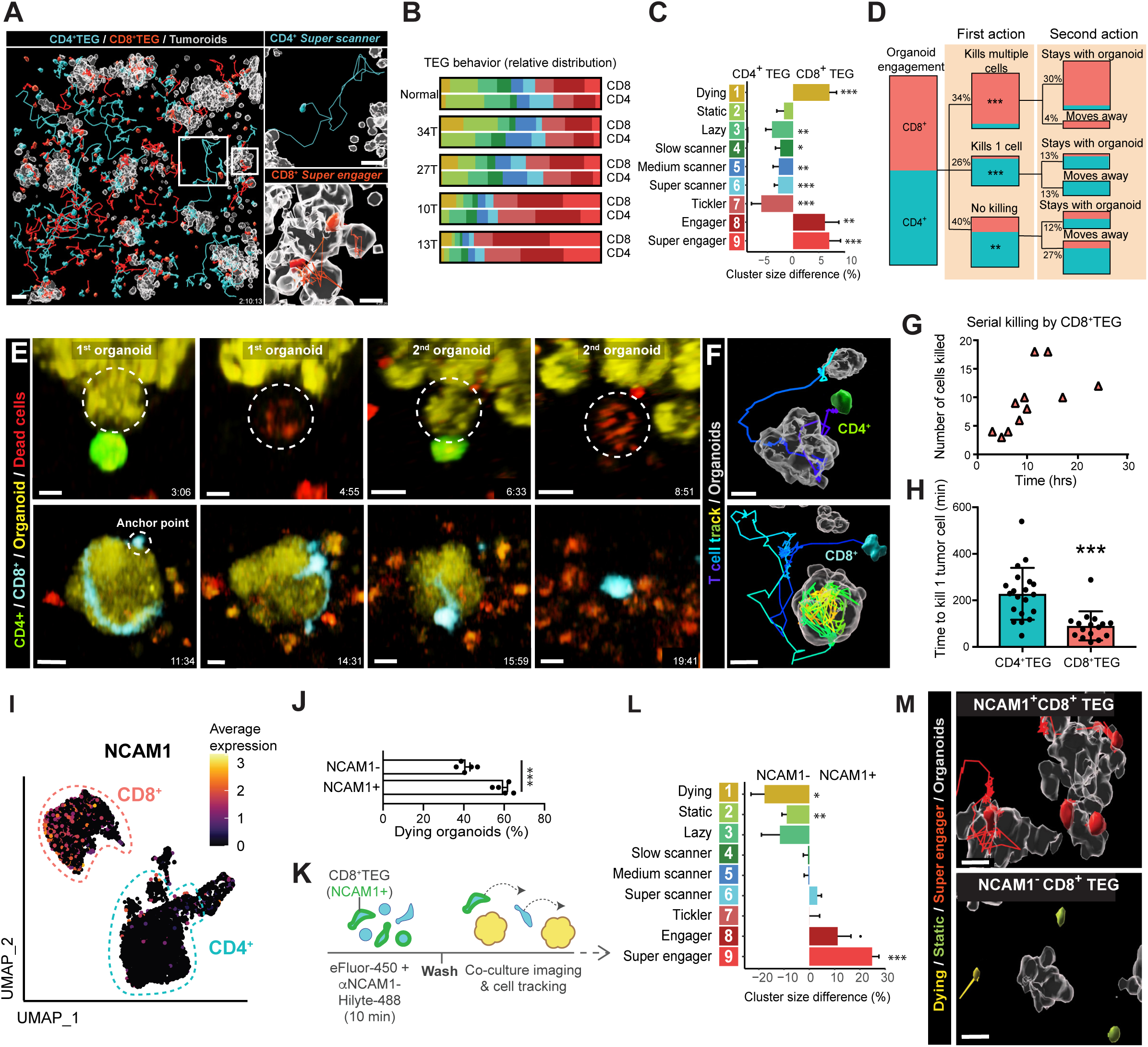
Unique targeting features of TEGs subpopulations and serial killer potential. (**A**) Representative 3D-rendered images of CD4^+^ (blue) and CD8^+^ (red) TEGs and their full tracks (up to 10 hrs) co-cultured with the 13T organoids (grey surface rendering at t = 0). Overview: Scale bar, 50 µm. Zoomed images: Scale bar, 30 µm (**B**) Relative behavioral cluster distribution of TEGs co-cultured with the indicated PDOs and a normal organoid culture. (**C**) Behavioral cluster size difference (%) between CD4^+^ and CD8^+^ TEGs co-cultured with the indicated PDOs and a normal organoid culture calculated from B. (n ≥ 3 independent experiments for each co-culture; mean ± s.e.m.) Linear regression model fitting with each well as a random effect: C9 p=7.52E-06; C8 p=0.0034; C7 p=0.00018; C6 p=0.000023; C5 p=0.0062; C4 p=0.01; C3 p= 0.001; C1 p=3.01E-06. (**D**) Quantification of the first action and second action of CD4^+^ and CD8^+^ TEGs after they engaged with an organoid. n = 3 replicates. Hypergeometric test was used to analyze cell type enrichment in each category. “Kills multiple cells” p<0.0001; “Kills one cell” p= 0.000015; “No killing” p= 0.0018. (**E**) 3D multispectral images showing a CD4^+^ TEG (green) that kills a 13T tumor cell (becomes red) in a first organoid (yellow) and a second tumor cell in a neighboring organoid (upper panel), and a CD8^+^ TEG (blue) killing a complete 13T organoid of ∼18 cells (yellow becoming red) in 11 hrs (lower panel). Scale bars, 30 µm. (**F**) Processed images of j showing 3D-rendered organoids (grey) at t = 0 and the CD4^+^ TEG (green) or the CD8^+^ TEG (blue) with their full track rainbow-colored for time. Scale bars, 10 µm. (**G**) Quantification of the number of cells killed in a sequence by CD8^+^ TEGs in time. (n = 3 independent experiments). (**H**) Quantification of the time it takes to kill one 13T tumor cell for CD4^+^ TEGs and CD8^+^ TEGs (n = 3 independent experiments). (**I**) UMAP embedding showing the expression levels of NCAM1. Color gradient represents the log_2_-transformed normalized counts of genes. (**J**) Quantification of the percentage of dying 13T organoids (% of total) at 10 hrs of co-culture with either sorted NCAM1^-^CD8^+^ TEGs or NCAM1^+^CD8^+^ TEGs (n = 5 independent experiments; mean ± s.e.m.). Two-tailed unpaired t test, p= 0.0001036. (**K**) Schematic representation of fluorescent labeling strategy of CD8^+^ TEGs with NCAM1 nanobody and efluor-450 to image and track NCAM1-positive versus -negative TEGs. (**L**) Behavioral cluster difference (%) between NCAM1^-^CD8^+^ TEGs or NCAM1^+^CD8^+^ TEGs co-cultured with 13T organoids. (n = 6 independent experiments; mean ± s.e.m.). Linear regression model fitting with each experimental replicate as a random effect: CL9 p=0.0002; CL8 p=0.07; CL2 p=0.005; CL1 p=0.02. (**M**) 3D-rendered images of 13T organoids (grey) from the same well with NCAM+ ‘super-engager’ CD8+ TEGs (top image) and NCAM- ‘lazy’ and ‘dying’ CD8+ TEGs (bottom image). Scale bars, 10 µm.

### NCAM1 associates with *super engager* behavior

Through single cell RNA sequencing (scRNAseq), we observed differential expression of NCAM1 in CD8^+^ TEGs (**Figures 3I, S3G and S3H; Table S3**). Although linked to cytotoxicity in both αβ and γδ T cells(Van Acker et al., 2017), this surface marker has not been examined in the context of cellular immunotherapy. We confirmed potent effector function related to NCAM1 expression, by showing that NCAM1^+^CD8^+^ TEGs had a higher capacity to kill 13T organoids compared to NCAM1^-^CD8^+^ TEGs (**Figure 3J**). To identify behavioral mechanisms underlying this high killing potential, we pre-labeled CD8^+^ TEGs with NCAM1 nanobodies (**Figure 3K)**, to directly compare NCAM1-positive and -negative populations within the same environment. NCAM1^+^CD8^+^ TEGs showed reduced *dying* and *static* behavior (**Figures 3L and 3M, and S3I**), supporting a higher *in vitro* persistence. Strikingly, NCAM1^+^CD8^+^ TEGs additionally showed a significant increase in *super engager* behavior compared to NCAM1^-^CD8^+^ TEGs (**Figure 3L and 3M**). Thus, surface marker expression can be linked to engineered T cell behavior, offering the opportunity to enrich for potent effector behaviors.

### Behavioral-transcriptomic profiling of TEGs

To generate insight into the transcriptional programs that underlie tumor-targeting dynamics revealed by BEHAV3D, we next performed single cell transcriptomic profiling of TEG populations enriched for different behavioral signatures, including a TEG population containing > 90% *super engagers* (**Figures 4A and 4B, and S4A; Video S1**). For each main TEG subset, effector CD8^+^ (CD8^+eff^), effector CD4^+^ (CD4^+eff^) and memory CD4^+^ (CD4^+mem^), profound transcriptional changes were observed upon 6 h co-culture with highly targeted 13T organoids, as compared to baseline (*no target control*) (**Figures 4C-4E**), showing that dynamic interplay with PDOs shapes the TEG transcriptomic profile. Behavioral probability mapping inferred from pseudotemporal ordering (**Figure S4B**) of the sequenced TEG populations (**Figure 4F**), revealed dynamic transcriptional programs that were highly conserved between CD8^+eff^, CD4^+eff^, and CD4^+mem^ TEGs (**Figure 4G**; Gene cluster (CL)1-3; 85% of genes; **Table S4**). These programs included genes to be down- (CL1) or up-regulated (CL3) by environmental stimuli or engagement to PDOs, as well as genes transiently expressed (CL2) along the pseudotime trajectory (**Figure 4G;** GO terms per CL in **Figure S4C**). This differential dynamic expression matched with known gene function, confirming robust ordering of TEGs, as shown by genes related to the CD3 signaling complex (LCK, SOS1, CD3E, CD3G; CL1; GO term ‘T cell activation’), known to be down-regulated upon T cell activation(Liu et al., 2000) in CL1 (**Figure 4H)**. NF-kB signaling, critical for tumor control(Barnes et al., 2015), and effector molecules including FASLG, IFNG, GZMB, TNF were found in CL3, with NF-kB signalling induced by environmental stimuli reaching maximum expression upon prolonged PDO-engagement, while the effector molecules appeared upon engagement (**Figure 4I**). In addition, CL3 contained genes related to rRNA processing that only increased upon prolonged engagement with organoids (**Figure 4H**), consistent with accelerated protein production in T cells following TCR engagement(Asmal et al., 2003; Tan et al., 2017). Finally, CL2 contained early activation markers CD69 and EGR1 with peak expression upon short organoid engagement, in line with IL-2 (CL3), known to be induced by EGR1(Collins et al., 2006), upregulated towards the end of the trajectory (**Figure 4I**). Thus, through our behavior-guided transcriptomics approach we robustly identified dynamic gene orchestration of TEG during tumor targeting.

**Figure 4.**
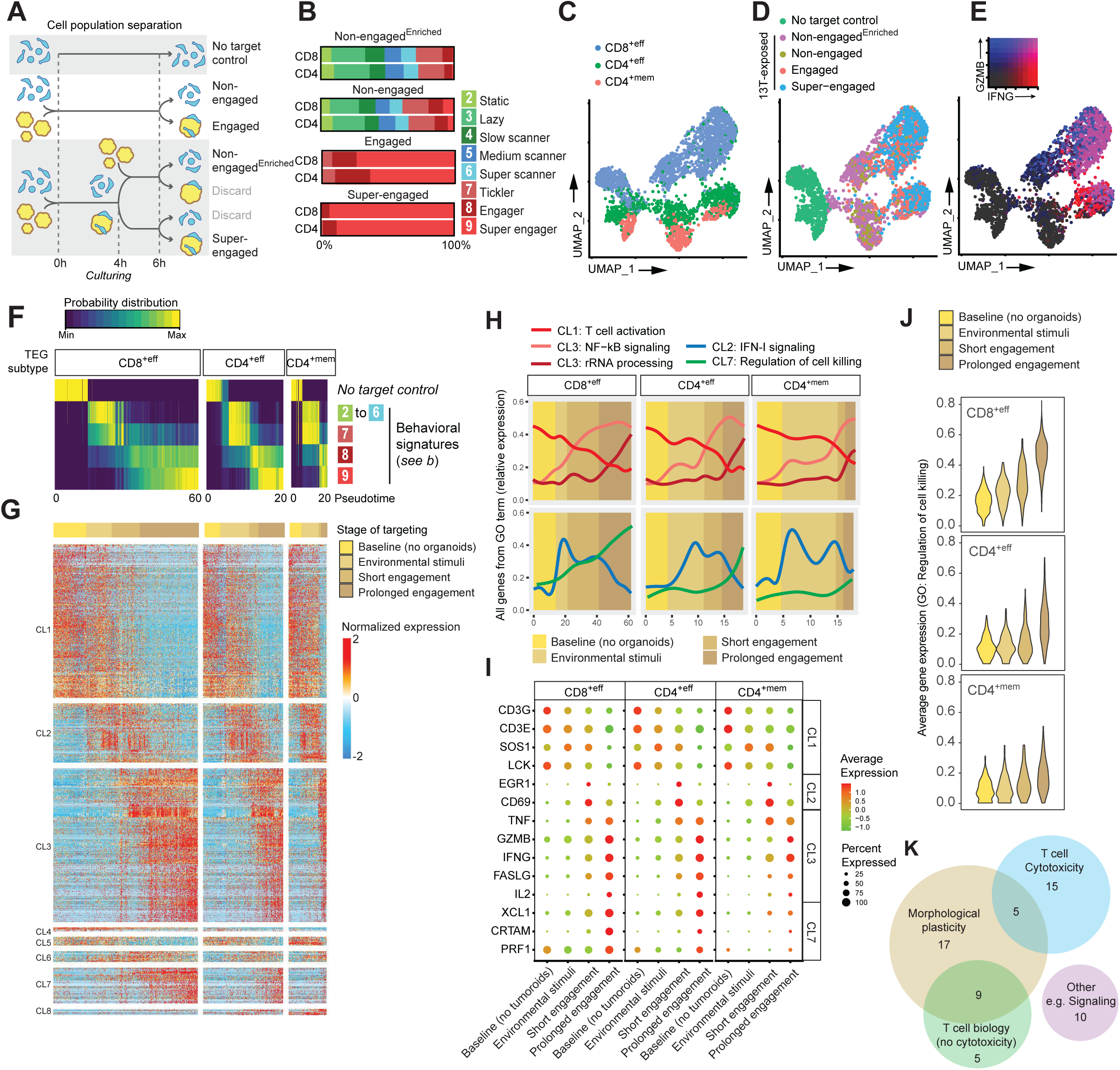
Behavioral-transcriptomic profiling of TEGs upon PDO exposure, engagement and killing. (**A**) Schematic representation of cell population separation for isolation and sequencing of *super-engaged*, *engaged*, *non-engaged*; *non-engaged^Enriched^*, and *no target control* TEGs. (**B**) Distribution of the 9 behavioral signatures described in Figures 2B and 2C of the indicated behavior-enriched TEG populations isolated at 6 hrs of co-culture. (**C-E**) UMAP embedding of pooled scRNAseq profiles showing distribution of, CD8^+eff^, CD4^+eff^, CD4^+mem^ TEGs (C), the 5 behavior-enriched TEG populations described in a (D), and normalized gene expression of IFNG and GZMB (E). Colors represent the log_2_-transformed normalized counts of genes. (**F**) Heatmap representing the probability distribution of different behavioral signatures and *no target control* over pseudotime for CD8^+eff^, CD4^+eff^ and CD4^+mem^ TEGs. Color represents the scaled probability for each behavioral group. (**G**) Heatmap showing normalized gene expression dynamics of TEGs upon exposure and engagement to 13T PDOs. Columns represent T cells ordered in pseudotime, rows represent the expression of genes, grouped based on similarity, resulting in 8 gene clusters (CL). CL1-3 represent gene expression patterns shared among TEG subsets. CL4-8 show different expression dynamics between TEG subsets. Horizontal color-bar (on top) represents the corresponding stage of targeting based on data in f. (**H**) Averaged gene expression over pseudotime for all genes from indicated GO terms for the indicated TEG subtypes. Graph background color-shading represent the corresponding stage of targeting. Line colors indicate GO term. (**I**) Gene-expression dot plot for a curated subset of genes at different stages of targeting. Rows depict genes. Dot color gradient indicates average expression, while size reflects the proportion of cells expressing a particular gene (%). (**J**) Violin plots for different TEG subtypes showing averaged expression of genes related to GO term ‘Regulation of cell killing’ enriched in CL7 from g. Colors indicate different stages of targeting. (**K**) Venn diagram depicting common and unique functions from 61 conserved genes composing a (serial) killer gene signature.

### Gene signature related to (serial) killing *super engager* TEGs

Of gene sets regulated in a TEG subset-specific manner (CL4-8; 15% of genes), CL7 contained genes mainly induced upon prolonged organoid engagement, including cytotoxic genes (e.g. PRF1, CRTAM, XCL1) (**Figures 4H and 4I;** GO: ‘Regulation of cell killing’). This cluster of genes was specifically induced in *super engager* CD8^+eff^, and to a lesser extent in CD4^+eff^ TEGs, and almost absent in CD4^+mem^ TEGs (**Figure 4J**), associating this gene cluster with potent (serial) killing T cells (**Figures 3D-3G**). Analysis of TEGs derived from a different healthy donor and co-cultured with another PDO culture (10T), confirmed that 61 out of the 83 genes of CL7 represent a conserved ‘killer’ gene signature (**Table S5**). Of these, we identified 20 genes related to T cell activation and cytotoxicity and 14 genes related to other T cell functions (**Figures 4K, and S4D**). However, importantly, we found 27 genes with no previously described T cell function (**Figures 4K, and S4D**). Overall, half of all conserved signature genes (31/61) and 17 out of the 27 novel genes were related to morphological plasticity processes, such as motility, cytoskeleton remodeling and adhesion (**Figure S4D**). Given that morphological plasticity is a key determinant of cell migration, many of the novel genes were found to have a role in promoting tumor cell migration and invasion, including ECM production and mesenchymal state induction (HEG1, BZW2, DCAF13, SQLE, PKIA). For some of these genes, such as CCT3 or AFAP1L2, the mechanism promoting migration is yet undescribed. In line with the prolonged organoid engagement behavioral feature of *super engager* TEGs (**Figure 2C**), we also found various genes related to cell adhesion, such as NCEH1, BYSL or EMP1. Finally, some genes had an additional function related to neurite outgrowth and dendritic pruning (SERPINE2, CHD4, NRTK1), potentially matching the long protrusion that were observed to occur in these serial killing TEGs **(Figures 3E, and Figure S3E and S3F; Video S1).** Thus, behavioral transcriptomics identified a specific gene signature induced in (serial) killing *super engager* TEGs.

### PDOs shape the dynamic gene signature of TEG during tumor targeting

To further explore our behavioral-guided transcriptomics approach, we next compared behavior-enriched TEG populations co-cultured between either highly sensitive 13T or intermediately targeted 10T PDOs. Distinct UMAP embedding of different TEG populations (**Figure 5A**) indicated that the patient-specific organoid exposure influences the dynamic TEG transcriptional profile. 41% or 61% of upregulated genes by environmental stimuli or upon prolonged PDO engagement in *super engagers*, respectively, were common between 10T- and 13T-co-cultured TEGs (**Figures 5B, and S5A and S5B**; **Table S6)**. Common *super engager*-related gene signatures included rRNA processing, NF-kB signaling and cytokine signaling (**Figure S5B**), and matched CL3 gene signatures (**Figure S4C**). However, 10T-co-cultured TEGs were characterized by induction of high cytokine expression upon prolonged PDO engagement, including TNF, IFNG and IL2, whereas IFN-I signaling genes were uniquely induced in TEGs co-cultured with highly sensitive 13T (**Figures 5C, and S5C**).

**Figure 5.**
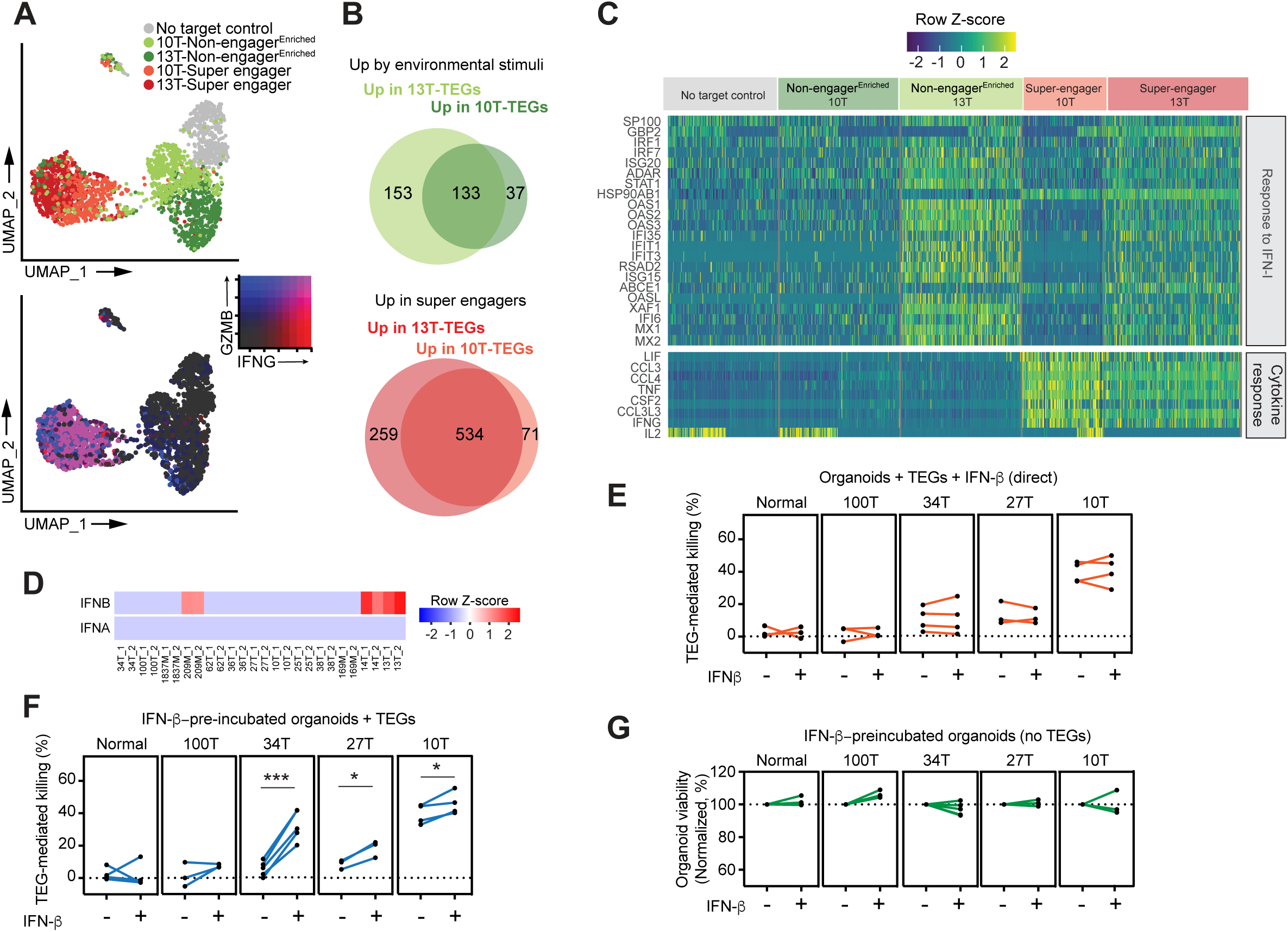
IFN-I signaling in PDOs primes TEG efficacy. (**A**) Top panel: UMAP embedding of pooled scRNA-seq profiles from *super-engaged* and *non-engaged^Enriched^* TEG populations co-cultured with either 13T or 10T PDOs; and *no target control* T cells. TEGs are colored per experimental condition. Bottom panel: UMAP plot showing the expression levels of IFNG and GZMB. Colors represent the log_2_-transformed normalized counts of genes. (**B**) Venn diagrams depicting common and unique genes upregulated in TEGs upon 13T and 10T organoid exposure (environmental stimuli; top panel) or prolonged engagement (*super engagers*; bottom panel). (**C**) Heatmap of gene expression for genes involved in functional annotations of interest (“Response to IFN-I”, “Cytokine response”), grouped according to TEG populations. (**D**) IFNA and IFNB expression in PDOs from the BC panel in Figure 1D. (**E-G**) Quantification of dying single organoids in presence or absence of recombinant IFN-β for the following conditions: organoids co-cultured with TEGs with direct addition of IFN-β, corrected for responses of LM1 control T cells (E); organoids pre-incubated with IFN-β for 24 hrs before co-culture with TEGs, corrected for responses of LM1 control T cells (F), and organoids pre-incubated with IFN-β for 24 hrs and cultured in absence of TEGs (G). Lines connect experimental replicates. (n≥3). Statistical analysis in (F) was performed by paired t test: 34T-IFN-β vs 34T-control p < 0.0006; 27T-IFN-β vs 27T-control p < 0.0216; 10T-IFN-β vs 10T-control p < 0.0402.

### IFN-β primes PDOs for TEG mediated killing

IFN-I signaling plays fundamental roles in anti-tumor immunity, yet with diverse and sometimes opposing functions reported for both tumor and immune cells, thereby making it difficult to fully comprehend and therapeutically exploit these effects(Boukhaled et al., 2021). IFN-I signaling was detected in 13T organoids (**Figure S1M**), which most prominently displayed increased RNA levels of the upstream mediator IFN-β, but not IFN-*α*, among our collection of PDOs (**Figure 5D)**. Secretion of IFN-β was confirmed by Luminex (**Figure S5D**), implying that IFN-β was the main mediator of IFN-I signaling observed in 13T. Interestingly, peak induction of IFN-I signaling in 13T-co-cultured TEGs was detected in non-organoid-engaging TEGs (from *static* to *super scanner* behavior), in line with a secreted source of IFN-β, while the pathway was shut down in *super engager* TEGs, suggesting a limited role of IFN-I signaling in direct killing behavior (**Figures 4F-4H**). Adding recombinant IFN-β to co-cultures of TEGs with various PDOs that showed low to medium sensitivity to TEG therapy (100T, 34T, 27T and 10T) indeed did not affect TEG targeting efficacy (**Figure 5E**). However, 34T, 27T and 10T organoids pre-treated with IFN-βshowed increased TEG-mediated killing, while IFN-β treatment did not impact organoid viability by itself (**Figures 5F and 5G**). These data support that IFN-β has limited impact on the killing capacity of *super engager* TEGs, confirming that dynamic IFN-I signaling is mainly associating with *static* to *scanner* behavior. Importantly however, IFN-I signaling increases the sensitivity of organoids to TEG therapy. Thus, behavior-guided TEG transcriptomics in relation to the type of organoid exposure reveals IFN-β to prime PDOs for targeting by TEGs. This illustrates the potential of the BEHAV3D to better understand and guide combinatory treatment approaches in a patient-specific manner.

## Discussion

Here, we provide an organoid-based 3D imaging-transcriptomic platform; BEHAV3D, for understanding the mode-of-action of cellular anti-cancer immunotherapies in a patient-specific manner. Using this pipeline, we report on the broad targeting potential of TEGs for breast cancer, poorly permissive to current immunotherapies (Esteva et al., 2019). In addition, by behavior-guided transcriptomics we have generated, to our knowledge, the first molecular map underlying the behavioral landscape of immune cells targeted to solid tumors. By exploiting these results, we were able to design an optimal sequence of IFN-I and TEG combination therapy to boost TEG organoid targeting.

Different from recent studies that have mapped the activation trajectories of murine immune cells during viral infection (Abbas et al., 2020), or human immune cells in normal physiology or cancer (Szabo et al., 2019), we here reconstructed activation trajectories for engineered T cells and uniquely exploited dynamic imaging data revealing their single-cell behavior. This allowed us to dissect gene programs induced by environmental stimuli, versus induction by short or prolonged tumor engagement, and thereby identify the gene signature of TEGs that (serially) killed tumor cells. This signature includes genes not previously linked to T cell function, thereby providing novel opportunities to potentially engineer next generation T cells with potent serial killing capability. Furthermore, multiple genes in this signature are associated with morphological plasticity. Such plasticity may underlie the remarkable cellular extensions of serial-killing TEGs, as observed in our 3D imaging data. Using these protrusions, TEGs intercalatebetween tumor cells while sequentially killing multiple tumor cells in the PDO, suggesting that morphological plasticity may be an important attribute in the targeting of solid tumors.

Type 1 IFNs have been described to be beneficial for the control of tumor growth, including in breast cancer, either by exerting direct antitumor effects (Dunn et al., 2005), or by improving the response to therapies, such as chemotherapy and checkpoint inhibition (Borden, 2019; Sistigu et al., 2014). Yet, opposite roles in inducing treatment resistance have been described as well (Benci et al., 2016; Boukhaled et al., 2021; Jacquelot et al., 2019). By using defined immune-organoid co-cultures, we have shown that an IFN-I signature intrinsic to tumor cells associates with TEG sensitivity, and that IFN-β primes tumor cells for more efficient targeting, rather than directly affecting TEG killing behavior. Thus, our data support the clinical use of IFN-I in combination with TEGs and possibly other cellular immunotherapies.

Adding to patient-specific drug responses observed in PDOs biobanks (Ganesh et al., 2019; Jacob et al., 2020; Ooft et al., 2019; Tiriac et al., 2018; Vlachogiannis et al., 2018; Yao et al., 2020), we show that not only killing efficacy, but also the underlying behavioral and molecular mechanisms of cellular immunotherapy differ between different PDO cultures. We even detected differences in killing dynamics between individual organoids belonging to the same PDO culture. This demonstrates that our platform captures the inter- and intra-patient heterogeneity, a major obstacle for treating solid tumors (Yamamoto et al., 2019). It is intriguing that gene signatures induced in TEGs upon organoid engagement were partly dictated by the type of PDO. In addition, the extent of IFN-βpre-treatment outcome on tumor targeting differed between PDOs, with the highly resistant culture 100T remaining unresponsive, whereas 34T displayed the highest (4-fold) increase in targeting. Together, these findings warrant caution regarding generalizing the outcome of immuno-oncology studies that use a single tumor model, and further supports the value of human organoid technology for development of personalized therapies.

Altogether, BEHAV3D combines organoid, imaging and sequencing technologies to offer a comprehensive platform that integrates multiple single-cell readouts, including tumor death dynamics, single-cell behavior and underlying transcriptomic profiling (**Video S1**). BEHAV3D may thus contribute to the efforts aimed at enhancing the efficacy of solid tumor-targeting by cellular therapies.

## Methods

### Human material

All human PDO samples were retrieved from a biobank through the Hubrecht Organoid Technology (HUB, www.hub4organoids.nl). Authorizations were obtained by the medical ethical committee of UMC Utrecht (METC UMCU) at request of the HUB in order to ensure compliance with the Dutch medical research involving human subjects’ act. The normal organoids were generated from milk obtained via the Moedermelkbank Amsterdam (Amsterdam UMC). Informed consent was obtained from all donors.

### Animal material

NOD.Cg-PrkdcscidIl2rgtm1Wjl/SzJ (NSG) mice purchased from Charles River Laboratories (France). Experiments were conducted in accordance with Institutional Guidelines under acquired permission from the local Ethical Committee and as per current Dutch laws on Animal Experimentation. Mice were housed in sterile conditions using an individually ventilated cage (IVC) system and fed with sterile food and water. Irradiated mice were given sterile water with antibiotic ciproxin for the duration of the experiment. Mice were randomized with equal distribution by age and initial weight measured on day 0 and divided into 10–15 mice per group.

### Organoid culture

Organoids were seeded in basement membrane extract (BME; Cultrex) in uncoated 12-well plates (Greiner Bio-one) and cultured as described previously(Dekkers et al., 2021; Sachs et al., 2017). Briefly, Advanced DMEM/F12 was supplemented with penicillin/streptomycin (pen/strep), 10 mM HEPES, Glutamax (adDMEM/F12+++), 1 x B27 (all from Thermo Fisher), 1.25 mM N-acetyl-L-cysteine (Sigma-Aldrich), 10 mM Nicotinamide (Sigma-Aldrich), 5 μM Y-27632 (Abmole), 5 nM Heregulin β-1 (Peprotech), 500 nM A83-01 (Tocris), 5 ng/ml EGF (Peprotech), 20 ng/ml FGF-10 (Peprotech), 10% Noggin-conditioned medium (NCM) (Cattaneo et al., 2020), 10% Rspo1-conditioned medium (RCM) (Broutier et al., 2016), and 0.1 mg/ml primocin (Thermo Fisher), and in addition with 1 μM SB202190 (Sigma-Aldrich) and 5 ng/ml FGF-7 (Peprotech) for PDO propagation (‘Type 1’ culture medium(Dekkers et al., 2021)), or with 20% Wnt3a-conditioned medium (WCM) (Broutier et al., 2016), 0.5 μg/ml hydrocortisone (Sigma-Aldrich), 100 μM β-estradiol (Sigma-Aldrich) and 10 mM forskolin (Sigma-Aldrich) for normal organoid propagation (‘Type 2’ culture medium (Dekkers et al., 2021)). Culture medium was refreshed every 2–3 days and organoids were passaged 1:2–1:6 every 7–21 days using TrypLE Express (Thermo Fisher). For co-culturing, organoids of a 5–12-day old culture (depending on PDO growth speed) were recovered from the BME by resuspension in TrypLE Express and collected adDMEM/F12+++. The organoid suspension was filtered through a 70 μm strainer (Greiner) to remove large organoids and pelleted before co-culturing. Organoids of passage 5 to 30 after cell isolation were used.

### Cell lines

Daudi (Gründer et al., 2012), HL60(Marcu-Malina et al., 2011) and Phoenix-Ampho cell lines were obtained from ATCC (authenticated by short tandem repeat profiling/karyotyping/isoenzyme analysis). Daudi and HL60 cells were cultured in RPMI media supplemented with 10% fetal calf serum (FCS) and 1% pen/strep (all from Thermo Fisher). Phoenix-Ampho cells were cultured in DMEM medium (Thermo Fisher) supplemented with 10% FCS and 1% pen/strep. All cells were passaged for a maximum of 2 months, after which new seed stocks were thawed for experimental use. Furthermore, all cell lines were routinely verified by growth rate, morphology, and/or flow cytometry and tested negative for mycoplasma using MycoAlert Mycoplasma Kit. Peripheral blood mononuclear cells (PBMCs) were obtained from Sanquin Blood bank (Amsterdam, The Netherlands) and isolated using Ficoll gradient centrifugation methods from buffy coats.

### Retroviral transduction of T cells

TEG001 (T cells engineered to express a highly tumor-reactive Vγ9Vδ2 TCR) (Gründer et al., 2012; Straetemans et al., 2015; 2018), LM1s (mock T cells engineered to express a mutant Vγ9/Vδ2 TCR with abrogated function) (Marcu-Malina et al., 2011), and TEG011 (mock T cells engineered to express HLA-A*24:02-restricted Vγ5/Vδ1 TCR; used as control for *in vivo* studies) (Kierkels et al., 2019; Scheper et al., 2013), were produced as previously described (Marcu-Malina et al., 2011). Briefly, packaging cells (Phoenix-Ampho) were transfected with helper constructs gag-pol (pHIT60), env (pCOLT-GALV) and pMP71 retroviral vectors containing both Vγ9/Vδ2 TCR chains separated by a ribosomal skipping T2A sequence, using FugeneHD reagent (Promega). Human PBMCs from healthy donors were pre-activated with anti CD3 (30 ng/mL; Orthoclone OKT3; Janssen-Cilag) and IL-2 (50 IU/mL; Proleukin, Novartis) and subsequently transduced twice with viral supernatant within 48 hrs in the presence of 50 IU/mL IL-2 and 6 mg/mL polybrene (Sigma-Aldrich). TCR-transduced T cells were expanded by stimulation with anti-CD3/CD28 Dynabeads (500,000 beads/10^6^ cells; Life Technologies) and IL-2 (50 IU/mL). Thereafter, TCR-transduced T cells were depleted of the non-engineered T cells.

### Depletion of non-engineered T cells

Depletion of non-engineered T cells was performed as previously described (Marcu-Malina et al., 2011). In short, αβ T cells were transduced with pMP71: γ-T2A-δ and incubated with a biotin-labelled anti-αβ TCR antibody (clone BW242/412; Miltenyi Biotec) followed by incubation with an anti-biotin antibody coupled to magnetic beads (anti-biotin MicroBeads; Miltenyi Biotec). Next, the cell suspension was applied onto an LD column and αβ TCR-positive (αβ TCR^+^) T cells were depleted by Magnetic-Activated Cell Sorting (MACS) according to the manufacture’s protocol (Miltenyi Biotec).

### Separation of CD4^+^ and CD8^+^ subsets of TEGs

In order to separate CD4^+^ and CD8^+^ TEGs and LM1s, we performed positive selection using either CD4 or CD8 Microbeads (Miltenyi Biotech) following manufacturer’s instructions. After incubation with magnetic microbeads cells were applied to LS columns and CD4^+^ or CD8^+^ TEGs or LM1s were selected by MACS. After MACS selection procedure, both Vγ9/Vδ2 TCR^+^ CD4^+^ or Vγ9/Vδ2 TCR^+^ CD8^+^ subsets of TEGs were separately expanded by using a rapid expansion protocol (REP) (Marcu-Malina et al., 2011) where TEGs were cultured in ‘TEG culture medium’ (RPMI-Glutamax supplemented with 2,5 – 10 % human serum (Sanquin), 1% pen/strep and 0.5M beta-2-mercaptoethanol) on a feeder cell mixture composing of irradiated allogenic PBMCs, Daudi and LCL-TM in the presence of IL2 (50 U/ml; Novartis Pharma), IL15 (5 ng/ml; R&D Systems) and PHA-L (1 μg/ml; Sigma-Aldrich). TEGs were stimulated biweekly by using the REP protocol. In order to monitor the purity of CD4^+^ and CD8^+^ TEGs, cells were analyzed by flow cytometry weekly prior to functional assays by using anti-pan γδ TCR-PE (Beckman Coulter), anti-αβTCR-FITC (eBioscience), anti-CD8-PerCP-Cy5.5 (Biolegend) and anti-CD4-APC (Biolegend) antibodies. TEGs with a purity lower than 90% were re-selected as described above. TEGs were used for co-culture assays 4 – 5 days after the last IL2/IL15/PHA-L stimulation.

### Sorting of NCAM1^-/+^ TEGs

CD8^+^ TEGs were harvested at day 8-10 of their REP cycle, stained in flow cytometry (FC) buffer (2% fetal bovine serum, 1x PBS) with Hilyte-488-conjugated NCAM1 nanobodies (1:400; QVQ) and LIVE/DEAD Fixable Near-IR Dead Cell Stain (1:1000; ThermoFisher) for 30 minutes at 4°C and consecutively sorted using a SONY SH800S or a FACS Aria Cell Sorter (BD Biosciences) into NCAM1^-^ and NCAM^+^ populations. Cells were rested for 16 h in ‘TEG culture medium’ and then used for co-culture.

### Live cell imaging of TEG and organoid co-cultures

LM1s or TEGs (20,000) were co-cultured with normal organoids, PDOs or control cell lines (Daudi or HL-60) in an effector to tumor cell (E:T) ratio of 1:3 or 1:25 (for **Figures 3D-3F**). CD4^+^ and CD8^+^ TEGs were mixed in a 1:1 ratio just before plating. Cells were incubated in 96-well glass-bottom SensoPlates (Greiner) in 200 µl ‘co-culture medium’: 50% ‘Type 1’ organoid culture medium, 50% ‘TEG assay medium’ (RPMI-Glutamax supplemented with 10% FCS and 1% pen/strep), 2.5% BME and pamidronate for the accumulation of the phosphoantigen IPP to stimulate tumor cell recognition(Marcu-Malina et al., 2011) (1:2000). ‘Co-culture medium’ was supplemented with both NucRed™ Dead 647 (2 drops per ml; Thermo Fisher) and TO-PRO-3 (1:3000; Thermo Fisher) for fluorescent labelling of living and dead cells (‘Imaging medium’). Combination of NucRed™ Dead 647 and TO-PRO-3 light up dead cell when excited with the 633 nm laser, and living cells when excited with the 561 nm laser (**Figures S1A and S1B**). Both were combined to achieve the most optimal fluorescent intensity ratio between dead and living cells for live cell imaging. Prior to co-culturing, TEGs were incubated with eBioscience™ Cell Proliferation Dye eFluor™ 450 (referred to as eFluor-450; 1:4000; Thermo Fisher) in PBS for 10 min at 37 °C to fluorescently label all TEGs. When CD4^+^ and CD8^+^TEGs were simultaneously imaged, eFluor-450, as well as Calcein AM (1:4000; Thermo Fisher) were used to label the different TEG subsets in PBS for 10 min at 37 °C. For NCAM1 pre-labeling experiments, a combination of eFluor-450 (1:4000; Thermo Fisher) and Hilyte-488-conjugated NCAM1 nanobodies (1:400; QVQ) was used to label CD8^+^ TEGs in PBS for 20 min at 37 °C prior to co-culturing. To prevent evaporation while imaging, 200 µl PBS was added to the wells surrounding the co-culture wells. The plate was placed in a LSM880 (Zeiss) microscope containing an incubation chamber (37 °C; 5% CO_2_) and incubated for 30 min to ensure settling of TEGs and organoids at the bottom of the well. The plate was imaged for up to 24 hrs with a Plan-Apochromat 20 x/0.8 NA dry objective with the following settings: online fingerprinting mode, bidirectional scanning, optimal Z-stack step size, Z-stack of 60 μm in total and time series with a 30 min (up to 60 conditions simultaneously; resolution 512 x 512) or 2 min interval (up to 4 or 10 conditions simultaneously; resolution 512 x 512 and 200 x 200 respectively). To minimize photobleaching of NCAM1-pre-labeled TEGs, the 488 nm laser was only activated 1 Z-stack each hr within the first hrs of imaging. Directly after imaging, the production of IFN-γ in the supernatant was quantitated using an ELISA-ready-go! Kit (eBioscience) and cell pellets were used to measure organoid viability using a CellTiter-Glo® Luminescent Cell Viability Assay (Promega).

### IFN-β stimulations

PDOs were harvested as described above and incubated in 96-well round bottom culture plates (Thermo Fisher) in 100 µl ‘Type 1’ organoid culture medium, supplemented with 2.5% BME, with or without the presence of 100 pg/ml recombinant human IFN-β (Peprotech). After 24 h incubation (37 °C; 5% CO_2_), TEGs or LM1s were added to either IFN-β-preincubated or unstimulated organoids (E:T ratio of 1:3) in 100 µl ‘TEG assay medium’, supplemented with 2.5% BME and pamidronate (1:1000), with or without the presence of 100 pg/ml recombinant human IFN-β (Peprotech). Medium without T cells was added for ‘organoid only’ controls. After 16 h incubation (37 °C; 5% CO_2_), plates were used to measure organoid viability using a CellTiter-Glo® Luminescent Cell Viability Assay (Promega).

### *In vivo* targeting by TEGs

Adult female NSG mice (15-16 weeks old) received sub-lethal total body irradiation (1,75 Gy) and subcutaneous implantation of an β-estradiol pellet (Innovative Research of America) on Day −1. On day 0, PDOs (1×10^6^ 13T organoid cells in 100 μl BME per mouse) were prepared as described previously (Dekkers et al., 2021) for subcutaneous injection on the right flank on Day 0, and received 2 injections of 10^7^ TEGs or TEG011 mock cells on day 1 and 6 in pamidronate (10 mg/kg body weight) as previously reported(Johanna et al., 2019). On day 1, together with the first T cell injection, all mice also received 0.6×10^6^ IU of IL-2 (Proleukin; Novartis) in IFA subcutaneously. Tumor volume was measured once a week using digital caliper and was calculated by the following formula: 0.4 x (length x width x width). Mice were monitored at least twice a week for weight loss and clinical appearance scoring (scoring parameter included hunched appearance, activity, fur texture, piloerection, respiratory/breathing problem). Humane endpoint (HEP) was reached when mice experienced a 20% weight loss from the initial weight, tumor volume reached 2 cm^3^, or when clinical appearance score of 2 was reached for individual parameter or an overall score of 4.

### Image processing

For 3D visualization, cell segmentation and extraction of statistics, time-lapse movies were processed with Imaris (Oxford Instruments), versions 9.2 to 9.5. The *Channel Arithmetics* Xtension was used for creating new channels to specifically identify organoids (live and dead) and eFluor-450-labeled or calcein AM-labeled TEGs (live and dead) and exclude cell debris. The Surface and ImarisTrack modules were used for object detection and automated tracking of both TEGs (autoregressive motion) and organoids (‘connected components’ or no tracking). The *Distance Transformation* Xtension was used to measure the distance between TEGs and organoids and thresholds for defining organoid–T cell interactions were visually determined. For tracked TEGs, time-lapse data containing the coordinates of each cell, the values of cell speed, mean square displacement, distance to organoids and dead cell dye channel intensity were exported. For experiments with NCAM1 pre-labelling, the mean intensities of the NCAM1 channel per T cell were exported. For tracked organoids, time-lapse data containing the coordinates of each organoid, the surface area, volume and mean dead cell dye channel intensity were exported.

### Imaging and sequencing data processing

Analysis of imaging and sequencing data was performed using R Studio version 4.0.2, as well as the following R packages: DESeq2, devtools, dplyr, dtwclust, eulerr, gganimate, ggplot2, ggpubr, ggrepel, gridExtra, hypergeo, kmodR, lme4, lmerTest, MESS, nlme, openxlsx, parallel, patternplot, pheatmap, plotly, plyr, png, purr, RColorBrewer, readxl, reshape2, rgeos, scales, Seurat, sp, spatstat, stats, tidyr, tidyverse, umap, VennDiagram, viridis, xlsx, zoo.

### PDOs killing dynamics

To quantify cell death dynamics of PDO cultures, > 5000 single organoids were analyzed at each time-point (48 time-points total). The mean dead cell dye intensity within single organoid surfaces, and values were rescaled to a range between 0 and 100 per experiment to normalize for variation in the absolute dead cell dye intensity. Per time point, organoids were classified as ‘dying’ when the mean dead cell dye intensity was above 7.

### T cell dynamics analysis and multivariate time series clustering

For the analysis of TEG behavior overtime, the following parameters were used: T cell death, contact with organoids, speed, square displacement and interaction with other T cells. Interactions between TEGs were measured by computing the distance to the nearest neighbor for each cell. To only include active interactions between TEGs that were not engaged to organoids, we considered cells whose mean speed for the last 20 minutes fell within the 3^rd^ quartile. A threshold for interacting T cells was defined as 10 µm between cell centroids and confirmed by visual inspection of imaging data. For each TEG time series, linear interpolation was used to estimate the values in few cases of missing time points. To compare time series independently of their length, cell tracks were cut to a length of 3.3 hrs. For each experimental replicate, the values of each of the numeric variables were converted to z-scores. To enhance the most discrepant values, only the 3^rd^ quartile range values were kept, while the rest was converted to the minimum values. Finally, each numeric variable was scaled to a range between 0 and 1 and normalized to the mean of the 0.99 quantile (per experimental replicate). For binary variables (TEG-TEG or TEG-organoid interactions), the values were labelled as 1 or 0 for interaction or no-interaction terms. Similarity between distinct cell tracks was measured using a strategy that allows for best alignment between time-series, previously applied for mitotic kinetics (Cai et al., 2018) or temporal module dynamics comparisons (Schafer et al., 2019). A cross-distance matrix based on the multivariate time-series data was computed using the dynamic time warping algorithm from the package “dtwclust”. To visualize distinct cell behaviors in 2 dimensions, dimensionality reduction on the multidimensional feature count table was performed by Uniform Manifold Approximation and Projection method (“umap” package) (Ali et al.; Becht et al., 2018). Clustering was performed using the k-means clustering algorithm with outlier detection. The identity of each cluster was defined by the mean time-series values of the different parameters (speed, square displacement, organoid contact, T cell interactions, cell death) within each cluster (**Figure 2C**). To confirm the identity of each cluster, T cell cluster assignments were back-projected to visualize the surfaces and tracks of particular T cell populations in the imaging dataset (**Figures 2A, S2A and S2B, and S3B**). A combination of datasets with distinct behavioral characteristics was used to construct a global TEG behavior atlas using the described methodology (**Figure 2B**).

### Cell behavior classification using a Random Forest classifier

For standardized integration of new experiments, we used a random forest classification approach (Breiman, L., 2001) in order to relate cell behavior to the nine behavioral signatures that we found in our global TEG behavior atlas (**Figure 2B**). To allow for inclusion of experiments with a low E:T ratio of 1:25, where the parameter of T cell interaction would be influenced as compared to the standard E:T ratio of 1:3, the following parameters were used: T cell death, organoid contact, speed, square displacement. The reference dataset used to build the global TEG behavior atlas was split into cell tracks to be used as a training dataset (95%) and a test dataset (5%). To reduce dimensionality, for each cell track, four time-series descriptive statistics were quantified and used to train the classifier. For numeric variables, the following measures were computed for each cell track: mean, median, the top 90% of the distribution, and the standard deviation. For binary values, such as the contact with organoids, the mean was calculated, as well as the mean and maximum of cumulative interaction. The random forest classifier was trained using 100 trees on the above-mentioned variables using the nine behavioral signatures as labels (**Figures S2C and S2D**). The test dataset was used to assess for accuracy of the classifier and to determine in which behavioral signatures the errors occurred (**Figure S2E**).

### Correlation between TEG behavior and organoid killing dynamics

To estimate the correlation between the onset of death in individual organoids and the engagement with T cells belonging to the engaging clusters (C7-9), we implemented a technique of sliding window correlation analysis, previously used for functional brain connectivity (Preti et al., 2017) and genome analysis (Burke et al., 2010). We calculated the Pearson correlation coefficient between the cumulative number of organoid contacts with TEGs from each cluster and the increase in dead cell dye intensity in each over a sliding window of 3 hrs (**Figures 2F and S2F**).

### T cell serial killing capacity analysis

For accurate long-term (up to 24 h) T cell tracking, TEGs were plated at a E:T ratio of 1:25. Tracks were manually corrected where required. Tracks were divided into shorter subtracks of 160 minutes. Using the random forest classifier described above, each subtrack was assigned to a behavioral signature (**Figures S2D and S2E**). The following statistics were calculated for each type of behavioral signature (9 clusters): for continuous variables (square displacement; speed, T cell death) the mean, median and standard deviation of the upper quantile were calculated, and for discrete variables (organoid contact and interaction with T cells) the mean, cumulative mean, maximum and cumulative maximum were calculated. Principal component (PC) analysis was used to reduce the dimensionality. The top 5 PCs were used to classify the change in behavioral signature over time (**Figures S3B and S3C**). Equivalent to the approach that was used for the full tracks in **Figure 2B**, we computed a cross-distance matrix based on the multivariate time-series data using the dynamic time warping algorithm and performed k-means clustering in UMAP space. The change in behavioral signatures was represented in a time-series color plot where each row represents one cell track and the color codes for behavioral signature (**Figure S3C)**. The relative proportion of CD4^+^ and CD8^+^ TEGs in each cluster was calculated and plotted next to each long-term classification (**Figure S3C**).

TEGs that engaged to organoids were back-projected to the imaging dataset and their first and second actions upon organoid engagement were visually analyzed (**Figure 3D**). TEG morphological plasticity was calculated by measuring the cell elongation (ratio between the longest and shortest axes) per cell and per individual timepoint. For each cell track, the plasticity was then computed as the ratio between the maximal and the minimal cell elongation (**Figure S3F**). The actions of CD4^+^ and CD8^+^ TEGs upon organoid engagement (**Figure 3D**) as well as the speed of killing and serial killing potential (**Figures 3G and 3H**) were quantified using Imaris software. Only TEGs were included for which tumor cell killing was clearly observed (usually visible as a decrease in living cell dye and an increase in dead cell dye, which co-occurred in many cases with target cell detachment from the organoid). For cases where a single organoid was fully killed by a single TEG, the number of cells killed by the TEG was calculated by dividing the killed volume by the average volume of a single 13T cell (2182 μm^3^). The killing rate of TEGs was measured as the time period from target cell engagement until tumor cell death (**Figure 3H**).

### NCAM1 pre-labelling quantification using 3D imaging data

Behavioral classification of NCAM1 pre-labelled TEGs was performed as described above, by predicting behavioral signatures with the Random Forest classifier. NCAM1^+/–^ TEGs were identified based on an NCAM1 intensity threshold in individual TEGs, visually defined at the timepoints where the 488 nm laser was turned on. To ensure inclusion of true NCAM1^-^ or NCAM1^+^ TEGs, two intensity thresholds were defined. Only tracks with a defined NCAM1^+^ or NCAM1^-^ identity were used for subsequent analysis. For each individual well, a difference in percentage of NCAM1^+^ and NCAM1^-^ TEGs was calculated per behavioral signature (**Figures 3L and S3I**).

### PDO bulk RNA sequencing

For bulk RNA sequencing characterization, RNA of PDOs grown in ‘Type 1’ culture medium was isolated according to the manufacturer’s protocol using the RNeasy Mini Kit (QIAGEN). Quality and quantity of the RNA samples and the libraries were measured with Agilent’s Bioanalyzer2100 and Invitrogen™ Qubit™ 3.0 Fluorometer. Quality control was done using FastQC, alignment has been done using STAR (https://github.com/alexdobin/STAR/releases/tag/STAR_2.4.2a) and reads have been mapped to the GRCh37 version of the human reference genome. Quality control on the bams was done using Picard. Read counts were generated with Htseq-count after which normalization is done using DESeq. RPKMs have been calculated with edgeR. For the library preparation the TruSeq Stranded mRNA Library Prep kit from Illumina was used. Sequencing was performed on the nextseq500 sequencer (also Illumina) with single-end 75bp reads. PDO cultures were ranked by responsiveness to TEGs (**Figure 1D**) and differentially expressed genes between the 6 most TEG-sensitive and 6 least TEG-sensitive cultures were analyzed. Genes exhibiting a more than 4-fold expression change with an adjusted p-value <0.05 after multiple hypothesis testing correction were used as input gene set enrichment analysis.

### SORTseq sample preparation

For sequencing of different behavior-enriched TEG populations (**Figure 4A**), TEGs (>0,8×10^6^ per condition) were either (1) co-cultured with 13T PDOs (E:T of 1:3) and separated into organoid-engaged (*engaged*) and organoid non-engaged (*non-engaged*) populations by 2 slow-spin (30 rcf) centrifugation steps at 6 h co-culture, (2) co-cultured with 10T or 13T PDOs (E:T of 1: 3) and separated at 4 hrs into organoid-engaged and organoid non-engaged populations by a slow-spin (30 rcf) centrifugation step, co-cultured for another 2h with or without addition of fresh PDOs, again followed 2 slow-spin (30 rcf) centrifugation steps to obtain *non-engaged^Enriched^* and *super-engaged* TEG populations, or (3) cultured for six hrs without addition of PDOs (*no target control*), using 12-wells culture plates (Thermo Fischer) and ‘co-culture medium’. To create single-cell suspensions, conditions containing organoids (all ‘*engaged’* TEG conditions) were treated with TrypLE for seven minutes at 37°C and washed with adDMEM/F12+++. Cells were then stained in FC buffer (2% FCS in PBS) with anti-CD3-APC conjugated antibodies (1:80; BioLegend) and LIVE/DEAD Fixable Near-IR Dead Cell Stain (1:1000; ThermoFisher) for 30 minutes at 4°C and sorted into 384-wells SORTseq plates using a FACS Aria Cell Sorter (BD Biosciences) and directly stored at −80°C until further processing.

### SORTseq library preparation and sequencing

All sorted plates were processed according to the CEL-Seq2 protocol with the total transcriptome amplification via poly-A RNA-capture, library preparation, and sequencing into Illumina sequencing libraries as previously described(Muraro et al., 2016). Paired-end sequencing (read1: 30 bp; read2: 120 bp) was used to sequence the prepared libraries using an Illumina NextSeq sequencer.

### Mapping and quantification of SORTseq data

SORTseq data were mapped and reads were counted, using STAR version 2.6.1a on the Hg38p10 human genome (annotated with GenCode v26). Plate-QC was performed using the Sharq pipeline (Candelli et al., 2018). Percentage of ERCC spike-in reads and mitochondrial mRNA reads versus total read count per cell were applied as QC parameters to identify systemic processing or pipetting errors over the plates. Cells with mitochondrial mRNA reads higher than 15%, ribosomal RNA content higher than 30%, or ERCC reads higher than 25% were excluded from the downstream analysis. Cells with fewer than 650 and higher than 4500 genes captured, and genes captured in fewer than 2 cells per plate were also excluded. After the initial QC steps, the ERCC spike-in reads were removed from the final count tables.

### SORTseq and 10x genomics data integration and TEG subpopulation analysis

For analysis of TEGs not exposed to organoids (**Figures 3I, and S3G and S3H**), 3 experimental replicates were used consisting of two datasets processed using SORT-seq and one dataset processed using 10x Genomics Chromium Single Cell 3′ gene-expression kit. SORTseq data was processed as described above. For the 10x dataset, (fresh, not co-cultured) TEGs were viability-enriched via FACS by DAPI staining (1:1000; Thermo Fischer) and loaded according to the standard protocol of the Chromium Single Cell 3′ Kit (v3). All the following steps were performed according to the standard manufacturer’s protocol. The library was sequenced on an Illumina Novaseq S1-flowcell and 19,000 reads/cell were collected. Single-cell RNAseq data were mapped, and counts of molecules per barcode were quantified using the cellranger(3.1.0) 10x software package to map sequencing data to the GRCh38(3.0.0) reference transcriptome supplied by 10x. Cells with mitochondrial mRNA reads higher than 15% and with fewer than 200 or more than 5000 distinct genes were excluded from the downstream analysis. Data were normalized by sequencing depth, scaled to 10,000 counts, log-transformed, and regressed against the UMI-counts and percentage of mitochondrial mRNA using the ScaleData function of the Seurat package. For integration of the 10x genomics (n = 1) and SORTseq (n = 2) datasets, we used previously published Seurat v3 data anchor-based integration(Stuart et al., 2019). Briefly, all three datasets were normalized using SCTtransform (Hafemeister and Satija, 2019) followed by selection of 5000 features for downstream integration. Transfer anchors were then learned and applied for integration of all datasets into a combined dataset. Cell visualization and placement in 2D view was achieved using principal component analysis (PCA) followed by Uniform Manifold Approximation and Projection (UMAP)(McInnes et al.). Shared nearest neighbor graph-based clustering was done using Seurat package’s FindNeighbors and FindClusters functions with a resolution of 0.8. For cell type identification marker genes for each cluster were calculated using the FindAllMarkers function and examined to profile marker genes that correspond to known cell types. Additional support for identifying cell subpopulations similitudes was achieved by analyzing the differentially expressed genes with a cell-type annotation tool(Cao et al.). Main marker genes used for TEG subpopulations identification are plotted in **Figure S3H.**

### Pseudotime Trajectory Inference

Two experimental SORTseq replicates of TEGs co-cultured with 13T PDOs, generated as described above, were used for trajectory interference (**Figure S4B)**. Proliferating T cells were excluded from the analysis since they did not show any dynamic inflammatory genes during the analysis. Afterwards, the gene expression table was log normalized with 10,000 scaling factor. Cell visualization and placement in 2D view was achieved using principal component analysis (PCA) followed by Uniform Manifold Approximation and Projection (UMAP) (McInnes et al., 2020). Shared nearest neighbor graph-based clustering was done using Seurat package’s FindNeighbors and FindClusters functions with the resolution of 2. Based on marker gene expression of CD8, CD4 and IL17RB (Terrier et al., 2010), TEGs were sub-clustered into 3 subtypes; IL17RB^-^CD8^+eff^, IL17RB^-^CD4^+eff^ and IL17RB^+^CD4^+mem^. Downstream analyses were done on each subset separately and compared with each other where mentioned. RunFastMNN function from SeuratWrappers package was utilized to correct for batch effects between the two SORTseq replicates. Unless specified, batch corrected UMAP values were used for visualization of single cells. We used Monocle3 (Cao et al., 2020) package to infer the pseudotime trajectory and significantly dynamic genes for each T cell subtype. For each cell subtype either *no target control* or *non-engaged^Enriched^* TEGs were designated as the root of the trajectory. In order to have comparable results from both Seurat and Monocle3 packages, the FastMNN batch corrected UMAP coordinates were imported and used throughout the trajectory analysis in Monocle3. In IL17RB^-^CD4^+eff^ and IL17RB^+^CD4^+mem^ subtypes, Monocle identified *no target control* cells as a separate partition. In order to have all cells along with a single pseudotime spectrum (e.g., not having several cells with a same pseudotime value), we added maximum pseudotime values of *no target control* T cells to pseudotime values of remaining cells in that subtype. For all TEG subtypes, significant dynamic genes along with the pseudotime trajectory were calculated and identified using Monocle3’s graph_test function using 1e-20 *q* value as the significance cutoff. Afterwards, using k-means clustering and also visual inspection of the genes’ behavior over the pseudotime, TEGs were clustered into sub-clusters with similar pattern (CL1-8; **Figure 4G**). The expression profile of the genes along with the pseudotime trajectory was plotted using pheatmap package(Kolde et al.) using row scaled (z-score) expression values. Smoothed gene(s) behavior was calculated and visualized recruiting gam smoothing function in ggplot2 package (Wilkinson, 2011).

### Behavior signature inference over the pseudotime

To align the pseudotime inference with the different behavioral signatures that we identified with BEHAV3D, we build a probability map distribution for different behavioral signatures over the pseudotime, based on the fundamental principle of transitivity of probabilistic distribution (**Figure 4F**). We defined three states of cells quantified by different methods:

- Behavioral_signatures (Bsig): {Static, Lazy, Medium-scanner, Scanner, Super scanner, Tickler, Engager, Super engager}. Behavioral signatures of cells identified by imaging (Figure 4B).
- Experimental_engagement_state (Expeng): {No target control, Non-engaged, Non-engagedenriched, Engaged, Super-engaged}. Cell distribution among different experimental conditions (Figure 4A)
- UMAP_cluster (Ucl): {1…X}. Cell assignment to distinct clusters grouping cells with similar gene expression. Shared nearest neighbor graph-based clustering was repeated several times using Seurat package’s FindNeighbors and FindClusters functions with a resolution ranging from 1 to 7.

From these three different cell states, the following information was quantified:

- p(Bsig|Expeng): For each Experimental_engagement_state we quantified the probability distribution of each Behavioral_signature (Figure 4F. This was achieved by reproducing the Experimental_engagement_states in silico on our imaging data. These values were calculated separately for CD4+ and CD8+ TEGs.
- p(Expeng|Ucl): For each UMAP_cluster, we quantified the probability of each Experimental_engagement_state to belong to this cluster.

Given these probabilities, we then quantified for each T cell the probability distribution of each unique Behavioral_signature in each UMAP_cluster, using the equation:

As a result, each cell was assigned a certain probability distribution for different behavioral signatures. To refine the probability map, the same process was repeated for 7 runs with different cluster sizes and the final probability distributions were averaged per cell. Note that for cells belonging to the *No target control* Experimental_engagement_state, a Behavioral_signature called *No target control* was assumed. The probability distribution along the pseudotime trajectory was plotted using pheatmap package^70^ of scaled values for each behavioral signature. Given that the non-engaged behavioral signatures (*Static, Lazy, Slow scanner, Medium scanner, Super scanner*) exhibited an identical probability map, their values were plotted together. For visualization purpose, extreme outlier values of skewed distributions were transformed to a maximal cutoff value. Based on the probability distribution of different behavioral signatures, the pseudotime was divided into 4 stages (Baseline (no organoids); Environmental stimuli, Short engagement, Prolonged engagement) for each TEG subtype (CD8^+eff^, CD4^+eff^ and CD4^+mem^).

### Differential gene expression analysis of TEGs co-cultured with distinct PDO cultures

For comparison of TEGs targeting 10T or 13T PDOs (**Figures 5A-5C**), SORTseq dataset was used including TEGs from distinct Experimental engagement states: *Non-engaged^Enriched^* and *super engager*. *No target control* TEGs were used as a control group. SORTseq data were mapped and quantified and visualized with UMAP as described above. Differential gene expression analysis was performed with the FindMarkers function from Seurat v3. Common and specific gene sets were filtered and visualized by Venn diagram with the VennDiagram package.

### Gene set enrichment analysis

The functional enrichment analysis in this study for pathway and biological processes annotations for gene sets of interest was conducted using ToppFun on the ToppGene Suite(Kaimal et al.) (**Figures 1H, S4C, and S5B**). An enrichment score was assigned based on gene enrichment ratio and log p value. For redundant annotations, the annotation with the highest gene enrichment ratio was selected. The pathways and biological processes with highest enrichment for gene set of interest were displayed in RStudio.

### Serial killer gene signature analysis

Genes of CL7 (**Figure 4G; Table S4 and S5**) were analysed to identify a unique signature for killer TEGs. 61/83 genes composing this cluster were common to TEGs incubated with 13T and 10T organoids and underwent extensive literature curation to identify genes with a known role in T cell cytotoxicity; T cell biology (not related to cytotoxicity); morphological plasticity or other processes such as GTPase signaling, ribogenesis and transcriptional regulation. Overlapping gene roles were plotted in a Venn diagram with the Venneuler package (**Figure 4K**).

### Statistical analysis

Statistical analysis was performed using R or Prism 7 software (GraphPad). Results are represented as mean ± s.e.m. unless indicated otherwise; *n* represents independent biological replicates. Two-tailed unpaired t-tests were performed between two groups, unless indicated otherwise. Pearson correlation was used for paired comparison between three different readouts (IFN-γ production, cell viability and live imaging). For live-cell imaging, the increase in dead cell dye between the first and last time point was used as measure. To compare tumor volume in mice treated with TEGs or TEG001 mock cells, two-way ANOVA with repeated measures was performed. To compare frequencies of different behavioral signatures between PDOs, a Pearson’s Chi-squared test was applied. To compare the percentage of dead organoids when TEGs were co-cultured with different PDOs, a one-way ANOVA followed by Bonferroni correction was performed. To estimate the change in correlation between 13T PDO death dynamics and cumulative contact with TEGs for different behavioral signatures, data was fitted to a linear mixed model with experimental replicate as random effect to account for variation between them. For cell type enrichment analysis of TEGs’ first and second action after engagement, a hypergeometric test was used (Fisher exact test). For comparisons of percentages of distinct TEG subtypes in the same well (CD4^+^ vs CD8^+^ or NCAM^+^ vs NCAM), for each behavioral signature data were fitted to a linear regression model with each individual replicate set as random effect to account for variation between them. For each fitted model, an analysis of variance was computed with an F-test. For comparison of IFN-β treatment, paired t test was performed.

## Data availability

RNA sequencing and imaging data is available upon request.

## Code availability

Upon request.

## Acknowledgements

We are grateful for the technical support from the Princess Máxima Center for Pediatric Oncology and the Hubrecht Institute and Zeiss for imaging support and collaborations. All the imaging was performed at the Princess Máxima Imaging Center. We thank the Princess Máxima Center Organoid Facility for organoid culture support, the flow cytometry facilities at the Princess Máxima Center and Laboratory of Translational Immunology (University Medical Center Utrecht) for cell sorting, the Princess Máxima Center Single Cell Genomics Facility for help with scRNAseq analysis, Single Cell Discoveries (https://www.scdiscoveries.com) for library preparations, and the Hartwig Medical Foundation (https://www.hartwigmedicalfoundation.nl/) for the sequencing. We also acknowledge the Hubrecht Organoid Technology (HUB) for providing PDOs, QVQ for providing NCAM1 nanobodies. We would like to thank Linde Meyaard, Jeffrey Beekman and the Dream3D^LAB^ for providing feedback on the manuscript, and Alberto Miranda-Bedate for pilot PDO sequencing analysis, Hannah Johnson for the voice over of the video and Anna Allemany for insightful advice for behavioral-transcriptomics inference. This work was financially supported by the Princess Máxima Center for Pediatric Oncology, Grants ZonMW 43400003, VIDI ZonMW 917.11.337, CRUK OPTIMISTICC C10674/A27140 (J.P., H.C), Netherlands Organ-on-Chip Initiative NWO 024.003.001 (J.P., H.C.), KWF Grants UU 2014-6790, UU 2015-7601, and UU 2019-12586 (J.K); Grant UU 2017-11393 (Z.S and J.K). K.K. was the recipient of a VENI grant from the Netherlands Organization for Scientific Research (NWO-ZonMW,016.166.140) and was a long-term fellow of the Human Frontier Science Program Organization (HFSPO, LT771/2015). A.C.R is supported by an ERC-starting grant 2018 project 804412. J.F.D was supported by a Marie Curie Global Fellowship and a VENI grant from the Netherlands Organization for Scientific Research (NWO).

## Author contributions

J.F.D grew organoids and performed imaging experiments with assistance from H.G.R., E.J.V. and M.G. M.A. designed and performed the computational analysis. M.A. and J.F.D. analyzed the data. M.B.B. and M.B.R. assisted with imaging data processing. J.F.D., A.K.L.W. and E.K. performed scRNAseq experiments. J.P. and D.F. processed PDO sequencing data. H.C.R.A. performed NCAM1 sorts. F.K. and P.B. analyzed the scRNAseq data. A.C. and E.K. produced TEGs and performed IFN-γ assays. H.G.R., I.J. and A.D.M. performed *in vivo* experiments. A.M.B assisted with computational analysis. R.L.I and M.B.R. made the video. O.K. provided organoid cultures. Y.B.E. and K.K. provided support and developed the co-culture live cell staining protocol that was used and further adapted. J.F.D, M.A. and A.C.R. designed the study and wrote the manuscript with support from A.M.M.E., E.J.W., H.G.S, Z.S., J.K. and H.C. This work was jointly supervised by A.C.R, Z.S, J.K, H.C who share senior authorship.

