## Supplemental information for "Behavioral-transcriptomic landscape of engineered T cells targeting human cancer organoids"

Dekkers, Alieva, *et al.*

**Supplemental information:**

Figures S1-5

Video S1

Tables S1-6

**Supplementary Figure S1. Related to Figure 1. Multi-spectral 3D imaging quantification of organoid killing and BC organoid biobank gene pathway signatures**. (**A**) Graph showing the emission spectra of the indicated fluorescent real-time cell dyes separately imaged by multispectral imaging with the lambda mode using the indicated lasers. Note that Nucred dead 647 and TO-PRO-3 have similar emission spectra. (**B**) Overview of the different fluorescent real-time cell dyes, the type of administration and which cell types they label in TEG-organoid co-cultures. (**C**) Schematic representation of the co-culture setup. Unlabeled organoids and eFluor-450-labeled TEGs were co-cultured in 'Imaging medium' in 96-well plates. (**D**) Quantification of death of individual PDOs in the presence of control TEGs expressing a mutated Vγ9/Vδ2 TCR (LM1s). Single organoids that crossed the mean dead cell dye intensity threshold of 7 (dashed lines) are considered dying (red lines). (**E**) Quantification of the percentage of dying single organoids (% of total) over time for each PDO co-cultured with LM1 control TEGs (n = 4 independent experiments; mean). (**F**) Quantification of death of individual PDOs in the presence of TEGs. Single organoids that crossed the mean dead cell dye intensity threshold of 7 (dashed lines) are considered dying (red lines). (**G-J**) Quantification of PDO targeting using a CellTiter-Glo® viability assay (G) or INFγ ELISA assay (I), upon 24 h co-culture of organoids with TEGs in the presence of pamidronate, and Pearson correlation plots between the outcomes of live cell imaging compared to CellTiter-Glo® measured viability (H) and INFγ ELISA (J). Data corrected for control LM1 T cell responses. (n = 4 independent experiments; mean ± s.e.m.). (**K-M**) Heatmap showing normalized gene expression (Row Z-score) for the indicated PDOs harvested at two different time points in culture (experimental replicates '_1' and '_2'). GO terms 'extracellular matrix (ECM) and ECM-associated proteins' (K), 'cytokines signaling in immune system' (M) and 'interferon signaling' (N) are presented, which were identified in the gene ontology enrichment analysis of differentially expressed genes between the six highest versus six lowest TEG-sensitive organoid cultures from **Figure 1H**.

**Supplementary Figure S2. Related to Figure 2. Properties of the 9 TEG behavioral clusters and random forest classification.** (**A,B**) Representative multispectral overview images (a; scale bars, 50 μm) and enlarged sections for clusters of interest (B; scale bars, 20 μm) of 13T organoids co-cultured with TEGs classified into 9 different behavioral clusters. (**C**) Schematic representation of the Random Forest classification pipeline and the resulting heatmap showing relative intensity values of T cell features indicated for each cluster resulting from the classification of the experiment in **Figure 2C**. (OC, organoid contact; Dis, square displacement; Sp, speed; TI, T-cell interaction; CD, cell death) (**D,E**) Error rate of the training data per cluster and overall for all trees (D) and correlation plot between ground truth cluster classification and predicted cluster classification (E). Color represents ground truth cluster. (**F**) Change in correlation between 10T organoid death dynamics (measured as increase in dead cell dye) and cumulative contact with TEGs (from behavior clusters 7-9). Data is represented as mean correlation per timepoint of all single organoids (n = 4 independent experiments).

Linear mixed model fitting with each experimental replicate as a random effect: C9 vs C8, $p < 2e-16$; C9 vs C7, $p < 2e-16$.

**Supplementary Figure S3. Related to Figure 3. Unique targeting features of $CD4^+$ and $CD8^+$ TEGs and behavioral signatures in relation to NCAM1 expression**. (**A**) Representative FACS plots showing CD4, CD8, $\alpha\beta$ TCR and $V\gamma9/V\delta2$ TCR expression for cultured $CD4^+$ and $CD8^+$ LM1s (mock) or TEGs. (**B**) Representative image of long-term tracking of TEGs in co-culture with 13T organoids (grey surface rendering at $t = 0$) showing full tracks (up to 20 hrs; rainbow-colored). Scale bar 50 μm. (**C**) Time series color plot showing long-term tracks of TEGs (co-cultured with 13T) and how they change their behavioral signature overtime for each time interval (0-3.3 hrs; 3.3-6.6 hrs; 6.6-10 hrs; 10-13.3 hrs). Colors indicate cluster identity for each TEG (see I). Long-term tracks were classified into 6 different groups that were named according to their most distinct long-term behavior and the proportion of $CD4^+$TEG and $CD8^+$TEG is indicated for each group. (**D**) Representative multispectral 3D-rendered images of a $CD4^+$ TEG moving away from a 13T organoid without killing. Scale bars, 20 μm. (**E**) Representative 3D multispectral images of TEGs co-cultured with 13T organoids showing 3 different $CD8^+$ TEGs with defined anchor points from which they scan and kill surrounding cells. Scale bars, 10 μm. (**F**) Quantification of fold increase in cell length indicating how much *super engager* TEGs can extend their shape as compared to their initial length. Each dot depicts individual cells pooled from 6 independent experiments. Boxplot depicts the median, first and third quartiles, whiskers extend 1.5 times from the interquartile range. (**G**) Uniform manifold approximation and projection (UMAP) plot shows TEGs unexposed to PDOs, pooled from three independent experiments. Colors indicate expression clusters identifying distinct cell subsets. (**H**) Gene-expression dot plot of a curated set of differentially expressed genes in each cell subpopulation. Rows depict cell subpopulations as in a, while columns depict genes. Dot color gradient indicates average expression, while size reflects the proportion of cells expressing a particular gene (**I**) Relative behavioral cluster distribution of NCAM1$^-$CD8$^+$ TEGs or NCAM1$^+$CD8$^+$ TEGs co-cultured with 13T PDOs.

**Supplementary Figure S4. Related to Figure 4. Behavior-guided transcriptomics of TEGs co-cultured with 13T organoids.** (**A**) Dynamic change of the percentage of TEGs exhibiting *super engager* behavior (C9) over time in co-culture. Color denotes TEGs co-cultured with 13T or 10T organoids and line type CD4$^+$ (dashed) or CD8$^+$ (solid) TEGs. The 6 hrs time point was selected for single cell TEG sequencing (dashed grey line). (**B**) Separate UMAP embeddings showing inferred pseudo-time trajectory of CD8$^{+eff}$, CD4$^{+eff}$, and CD4$^{+mem}$ TEGs. Color scale represents the inferred pseudotime. (**C**) Functional enrichment analysis for biological processes and pathways from gene clusters (CL) that are downregulated (CL1), upregulated (CL3), or transiently expressed (CL2) over the pseudotime trajectory of TEGs targeting 13T organoids. CL1-3 are represented in **Figure 4G**. (**D**) Gene-expression dot plot of the 61 conserved genes composing the (serial) killer gene signature separated by function. Rows depict genes, while columns depict stage of targeting. Dot color gradient indicates average expression, while size reflects the proportion of cells expressing a particular gene.

**Supplementary Figure S5. Related to Figure 5. Behavior-guided transcriptomics of TEGs co-cultured with 13T and 10T organoids.** (**A**) Heatmap showing normalized gene expression of behavior-enriched TEG populations co-cultured with 10T or 13T organoids, or cultured without PDOs (*No target control*). Columns represent cells ordered by TEG populations and rows represent the expression of genes. Shown are 534 genes induced upon prolonged organoid engagement (*super engagers*) in both 10T and 13T co-cultures from **Figure 5B**. (**B**) Functional enrichment analysis (conserved biological processes and pathways) of genes induced in both 10T- and 13T-co-cultured *super engager* TEGs (shown in A). (**C**) Functional enrichment analysis of genes differentially expressed between 10T- and 13T-co-cultured *super engager* TEGs. Top differentially regulated biological processes and pathways are shown. (**D**) IFN-β concentration measured for the different organoid cultures in **Figure 5 E-G**.

**Video S1.** Visual summary of the technology and main findings of the article.

**Table S1.** Characteristics of organoid cultures derived from 14 breast cancer patients.

**Table S2.** DEG analyses between the sixth highest versus lowest TEG- sensitive tumoroid cultures from **Figure 1D**.

**Table S3.** DEG analyses between different TEG subpopulation identified in **Figure S5A**.

**Table S4.** List of genes clustered based on distinct expression dynamics in TEGs during 13T PDO exposure and targeting corresponding to **Figure 4G**.

**Table S5.** Conserved genes of the (serial) killer TEG signature corresponding to **Figure 4K**. Genes with known or previously undescribed T cell function are highlighted in blue or green, respectively.

**Table S6.** DEG analyses and common genes corresponding to **Figure 5** analyses.
