## Supplemental Figures for "Behavioral-transcriptomic landscape of engineered T cells targeting human cancer organoids"

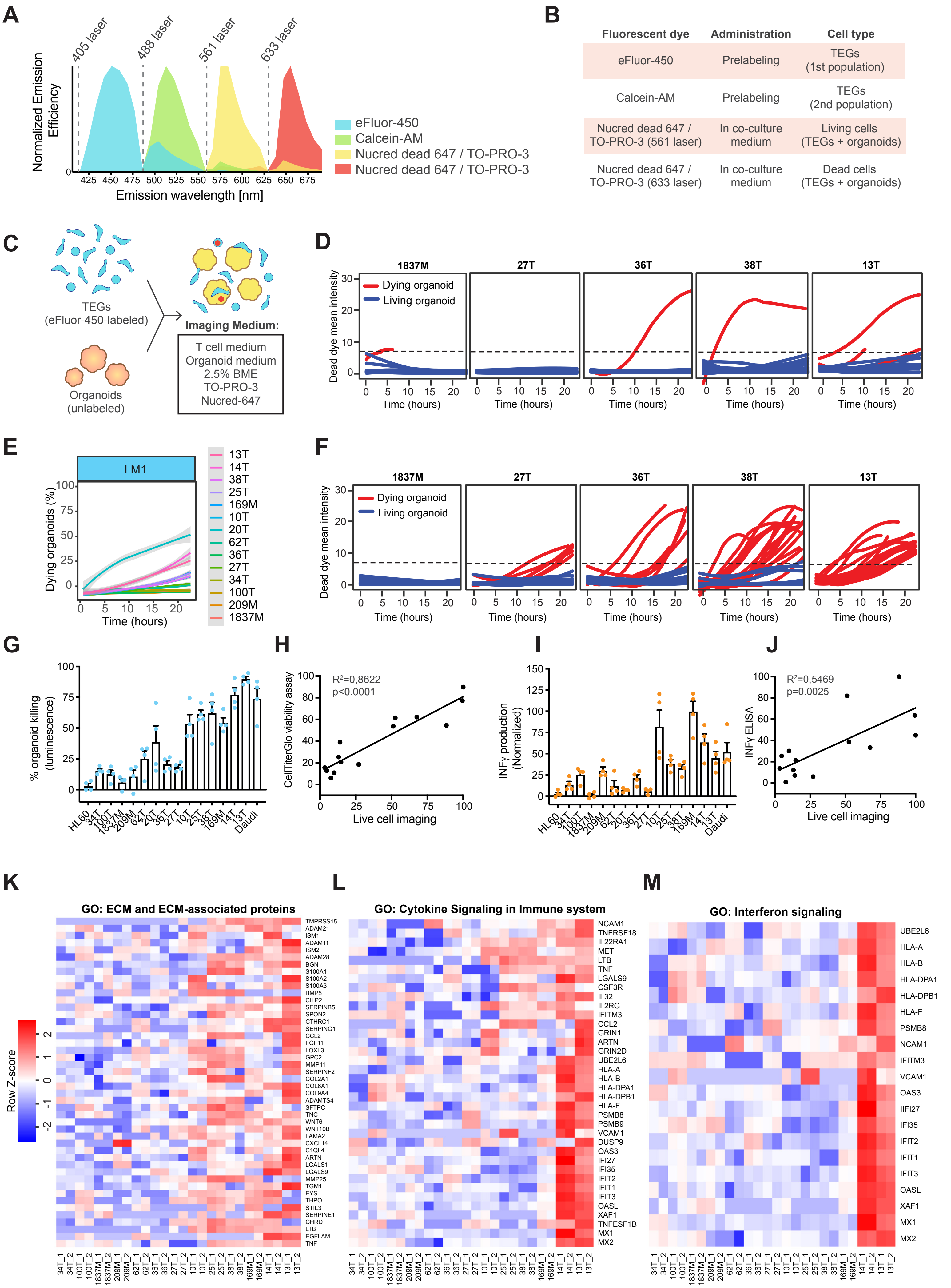


**Figure S1. Related to Figure 1**

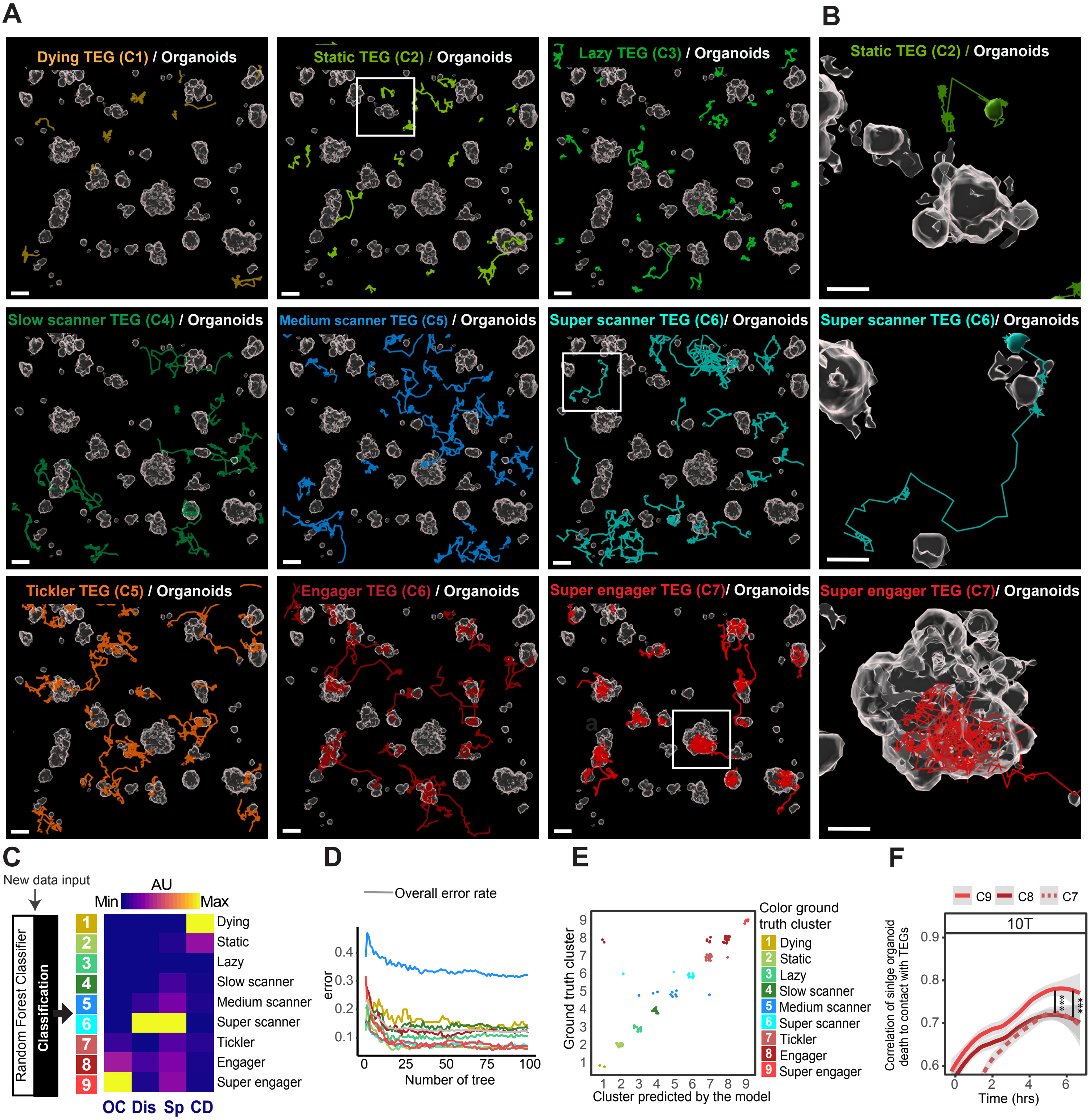


Figure S2. Relate to Figure 2

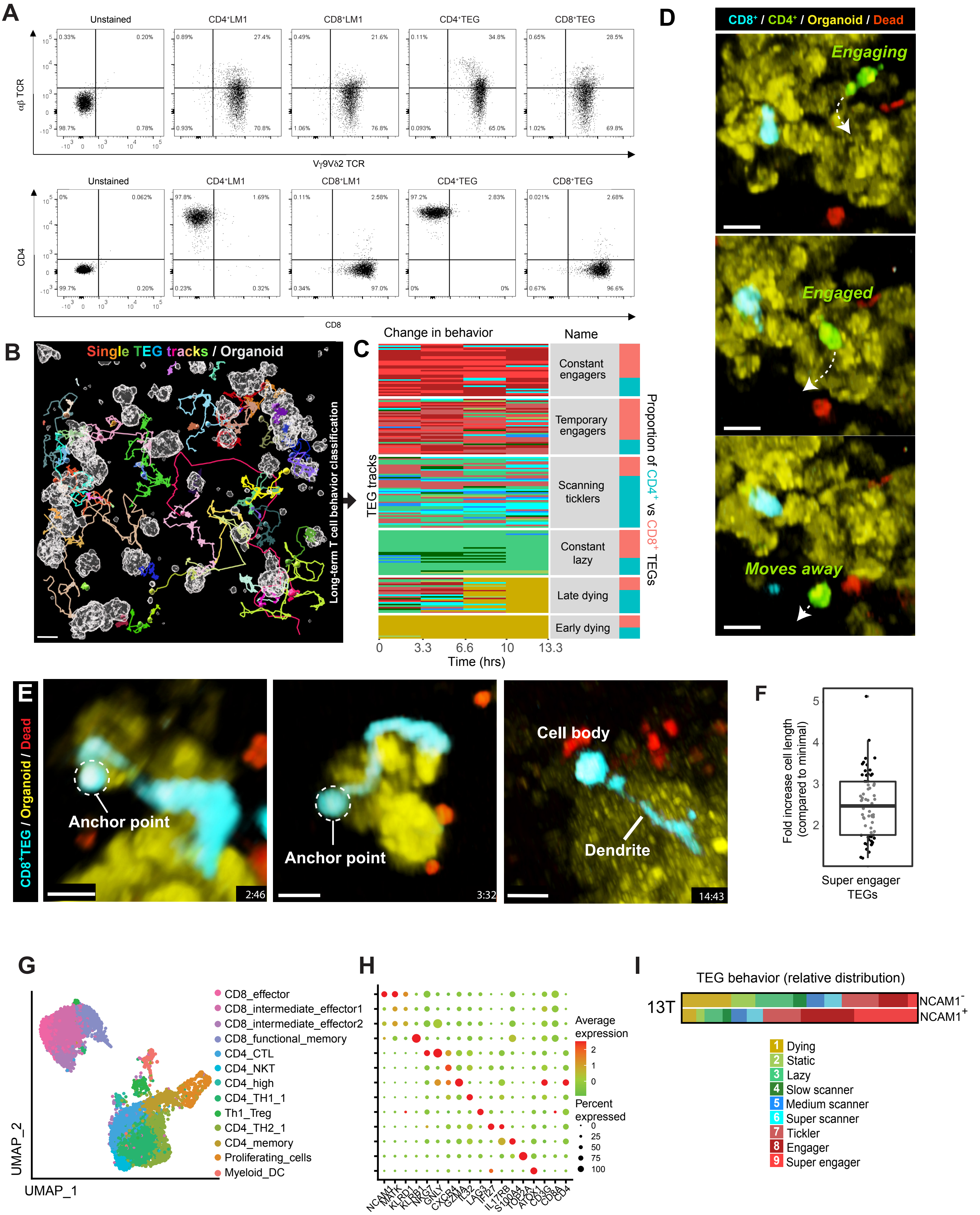


Figure S3. Related to Figure 3

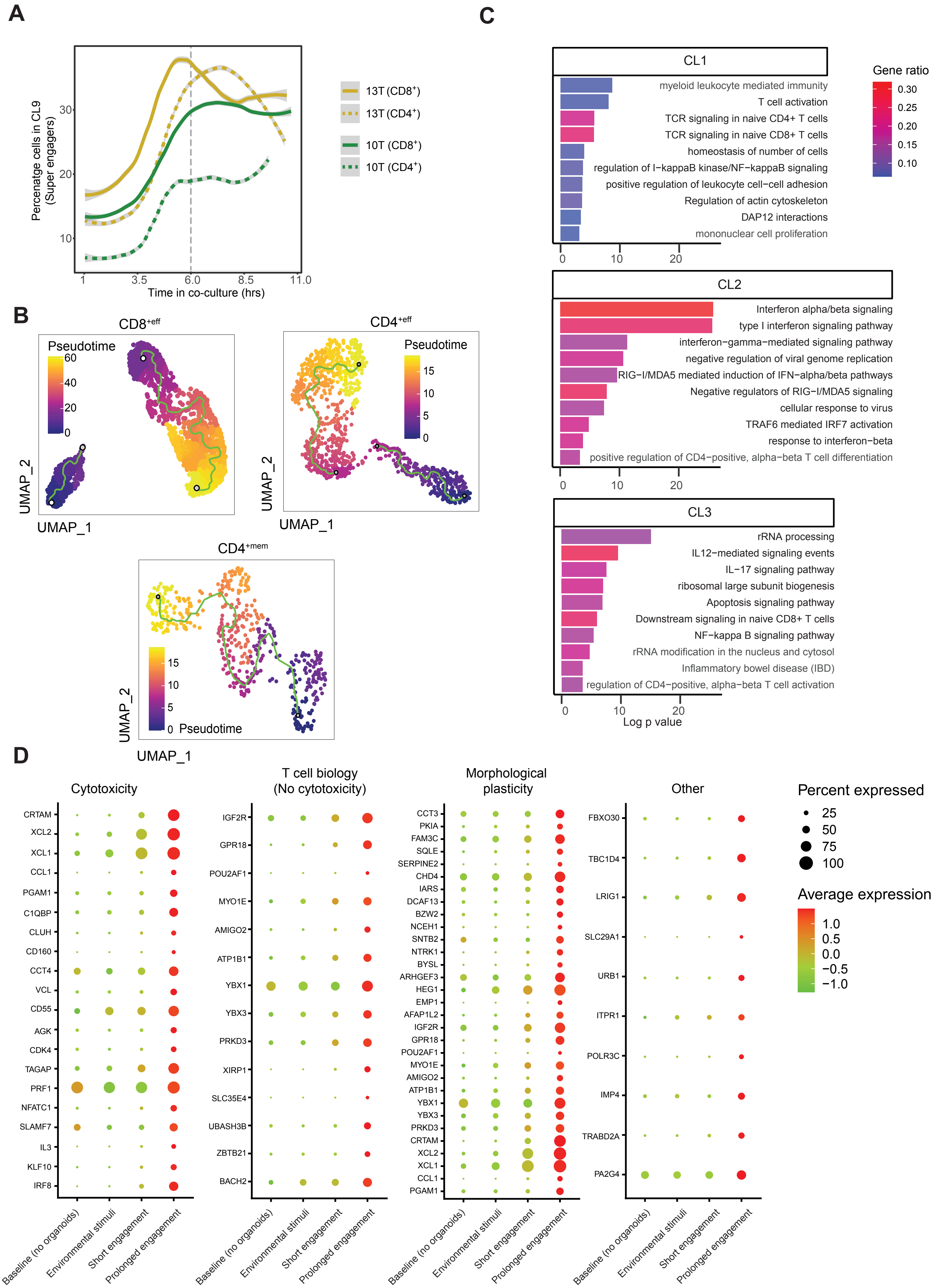


**Figure S4. Related to Figure 4**

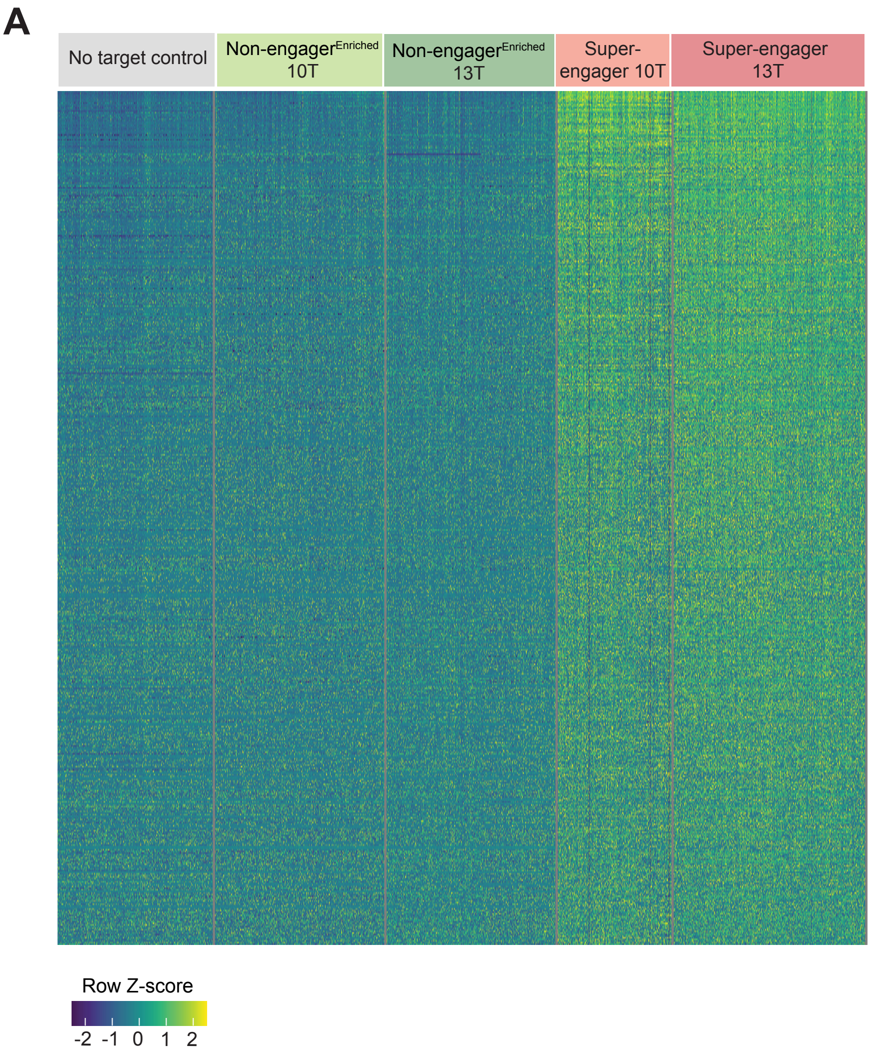


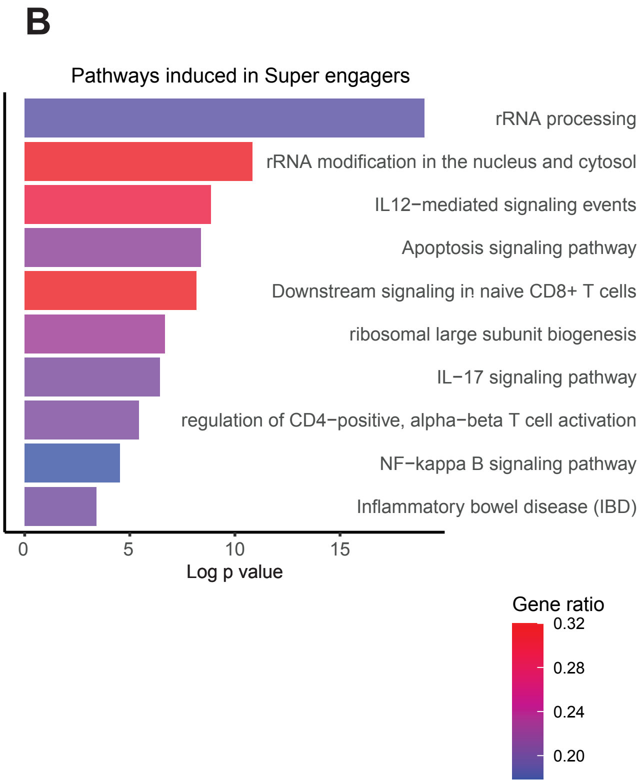


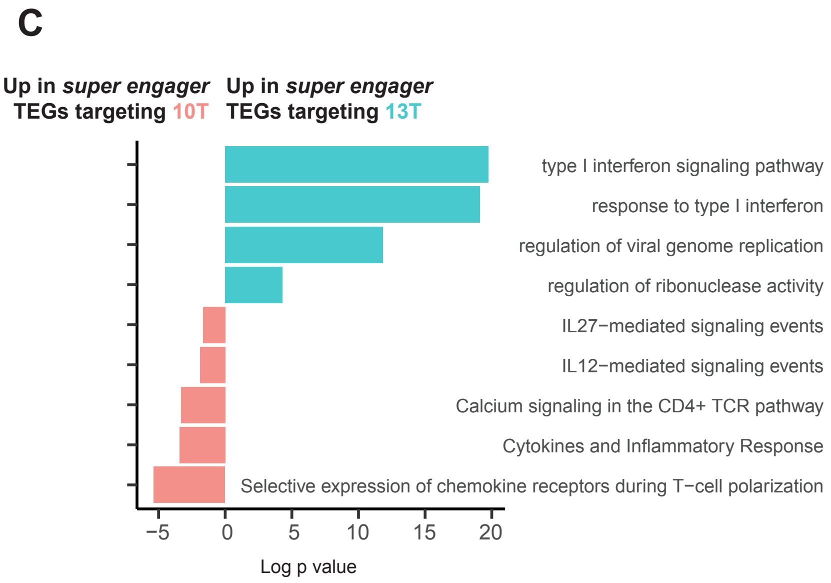


**D**

| Organoid | IFN-β (U/ml) |
|---|---|
| Normal | 0,94* |
| 100T | 0,99* |
| 34T | 0,89* |
| 27T | 1,42* |
| 10T | 2,59* |
| 13T | **6,47** |

*Value below standard range

**Figure S5. Related to Figure 5**
